# Seasonal Rebalancing of Assembly Processes and Cross-Domain Network Restructuring in Microbial Communities of a Large Eutrophic Lake

**DOI:** 10.64898/2026.07.29.741506

**Authors:** Shanmei Zou, Lydia Smith, Xinke Yu, Tiantian Sun, Xuemin Wu, Beixin Wang, Tyler Linderoth, Sven Katzenmeier, Thorsten Stoeck

## Abstract

Harmful algal bloom (HAB) succession in eutrophic lakes arises from seasonal rebalancing of selection, dispersal, and drift across interacting microbial domains, yet process attribution remains limited due to the scarcity of cross-domain analyses. Using multi-marker eDNA metabarcoding (16S rRNA, 18S rRNA, 23S rRNA), we show that bacterioplankton, cyanobacteria, phytoplankton, and zooplankton in Lake Taihu undergo coordinated spring–summer transitions, with inter-seasonal turnover far exceeding spatial heterogeneity. Assembly regimes shifted coherently across domains: spring communities were dominated by strong homogeneous selection—especially among cyanobacteria and microeukaryotes—whereas summer communities moved toward drift- or neutrality-like dynamics under bloom-stage mixing that enhanced dispersal. Null- and neutral-model diagnostics supported these trends. Cross-domain co-occurrence networks restructured seasonally, with summer networks becoming more connected and substantially more robust, indicating that bloom conditions foster cohesive, robust interaction structures rather than destabilization. These findings provide evidence that bloom progression restructures aquatic microbiomes through a seasonal rebalance of selection, drift, and dispersal coupled with adaptive strengthening of cross-domain connectivity, providing a process-explicit framework for understanding community stability and ecosystem resilience in eutrophic lakes.

## Introduction

Harmful algal blooms (HABs) are major disturbances in freshwater ecosystems, threatening biodiversity, restructuring food webs, and degrading water quality through toxins, hypoxia, and odor problems [1, 2]. Although nutrient enrichment and climate warming are recognized as primary drivers, bloom development in eutrophic lakes emerges from microbial consortia composed of bacterioplankton, viruses, cyanobacteria, and microeukaryotes (phytoplankton and zooplankton). These consortia mediate nutrient cycling, organic-matter processing, and top-down control, and reorganize seasonally in response to physical forcing and resource shifts [3–5]. Understanding bloom dynamics therefore requires resolving how microbial communities reassemble over time. Long-term lake studies—from Mendota to Erie [6–9]—consistently show seasonally structured bloom patterns driven by physical– chemical–biological feedbacks.

Recent work across lakes such as Mendota, Donghu, and peri-Alpine systems has begun to resolve cross-domain microbial dynamics and network responses to seasonal and physical forcing [6, 10–12]. Two-decade metagenomics from Lake Mendota revealed that invasive species can dramatically restructure protistan communities, driving declines in cryptophytes and increases in alveolates and diatoms [6]. Such community shifts are underpinned by network architecture: nine-year analyses in Lake Donghu showed that microbial networks are most stable in autumn, with keystone taxa (e.g., Burkholderiales) playing critical roles in maintaining seasonal stability [10]. The drivers of these patterns extend beyond local processes—century-long sedimentary DNA from peri-Alpine lakes demonstrated that cyanobacterial communities exhibit weak dispersal limitation at regional scales, with climate warming (rather than nutrients) emerging as the dominant assembly driver, selecting for traits such as buoyancy regulation [11]. Microbial communities reflect deterministic (selection, environmental filtering) and stochastic (drift, dispersal) forces, whose relative contributions shift as seasonal changes alter temperature, stratification, light, and nutrients [13–16]. For instance, warming and stratification strengthen deterministic selection [15, 16], while spring mixing enhances dispersal and stochastic assembly; light further reinforces deterministic filtering by selecting photosynthetic traits like cyanobacterial buoyancy [12]. These shifts leave characteristic signatures in β-diversity, neutrality deviations, and seasonal restructuring of cross-domain networks that regulate bloom dynamics [17–23].

Despite this progress, process-based understanding of HAB succession remains limited for three reasons. First, most studies analyze only one of bacteria, cyanobacteria, and microeukaryotes separately, limiting insight into coordinated cross-domain assembly [21, 22, 24, 25]. Second, although network and neutral-theory tools are widely applied [26–28], few studies integrate β-diversity, null, and neutral models to partition selection, dispersal, and drift across domains. Co-occurrence networks reveal non-random associations but cannot alone distinguish ecological processes or turnover sources [17, 18, 20, 22, 27, 29, 30]. Third, the mechanistic link between seasonal network restructuring and network robustness-defined as the resistance of co-occurrence networks to simulated species removal remains poorly understood. Integrating these complementary approaches is essential because each addresses a distinct facet of community dynamics: β-diversity partitioning reveals whether compositional changes are driven by species replacement or abundance shifts [17, 18]; null and neutral models (Beta Nearest Taxon Index-βNTI, Raup-Crick metric based on Bray-Curtis dissimilarity-RC_bray, Sloan neutral model) disentangle the relative contributions of deterministic selection, dispersal, and ecological drift [19, 20, 26, 27]; and network analysis uncovers how cross-domain interactions restructure in response to environmental forcing [21, 22, 24]. Individually, each method provides only partial insights; together, they enable a holistic, process-explicit understanding of how microbial communities assemble, respond to seasonal forcing, and maintain stability during bloom transitions. Although recent studies indicate enhanced network stability under strong seasonal forcing and show that microscale heterogeneity sustains coexistence in mixed layers [31, 32], we still lack an integrated framework to evaluate how such structural reorganization contributes to ecosystem stability during bloom events [33–35]. Addressing these gaps requires an integrated framework that explicitly links microbial community composition to the underlying ecological processes— selection, dispersal, and drift—while unifying cross-domain planktonic groups to resolve community reassembly and cross-domain interaction network restructuring across seasonal transitions. Such integration is increasingly critical, as multi-year observations from lakes such as Mendota and Erie show that microbial networks and bloom resilience arise from interactions between deterministic forcing (temperature, hydrodynamics) and stochastic dispersal and drift [6–8, 36]. At finer spatial scales, trait-based and microscale studies [12, 31, 32] further demonstrate that turbulence and chemical gradients restructure microbial associations and promote coexistence, underscoring the need for cross-domain, process-explicit analyses that incorporate physical, biological, and stochastic drivers.

Recent advances in multi-marker environmental DNA (eDNA) metabarcoding now make this feasible by simultaneously profiling bacterioplankton, cyanobacteria, and eukaryotic plankton through 16S rRNA, 18S rRNA, and 23S rRNA gene markers [37–39]. This unified eDNA framework enables mechanistic dissection of how deterministic and stochastic processes jointly govern microbial reassembly and cross-domain coupling throughout HAB succession.

Lake Taihu—China’s third-largest freshwater lake—is one of the world’s most intensively studied eutrophic systems and a key counterpart to long-term bloom-observing lakes such as Mendota and Erie. This shallow, polymictic lake has experienced increasingly severe cyanobacterial blooms since the early 2000s [40–42]. The spring-to-summer transition marks a shift from early proliferation to peak biomass, accompanied by pronounced changes in temperature, stratification, nutrient dynamics, and community structure. In 2017, an extreme bloom covering over 60% of the lake surface provided a natural large-scale experiment to test how strong bloom pressure and seasonal forcing reshape the balance among selection, dispersal, and drift, driving turnover-dominated reassembly and cross-domain network reconfiguration [43]. Spatial analyses of this event revealed pronounced heterogeneity in bloom occurrence across the lake, with blooms typically initiating in the northwestern region and subsequently spreading to the central basin [43]. This setting offers a unique opportunity to extend process-explicit frameworks from other lake systems into a multi-marker, cross-domain context for directly comparing deterministic and stochastic mechanisms of bloom succession [6, 11, 26].

In this study, we conducted the first large-scale, multi-marker eDNA survey of bacterioplankton, cyanobacteria, eukaryotic phytoplankton, and zooplankton across Lake Taihu during spring (initial bloom) and summer (peak bloom) in 2017. We asked a mechanistic question: how does the spring-summer transition rebalance the fundamental forces of selection, dispersal, and drift to drive community reassembly and bloom succession across multiple microbial groups? To address this, we designed an integrative analytical framework structured around three sequential questions:

First, we sought to quantify the magnitude and nature of community change. We asked: (i) Is the spring-summer transition characterized primarily by species turnover (i.e., replacement) or by shifts in relative abundances? And does this temporal signal overwhelm spatial heterogeneity? To answer this, we combined β-diversity partitioning (to distinguish turnover from abundance gradients) with PERMANOVA (to compare the effect sizes of season vs. space). We further quantified distance-decay relationships to assess how community similarity declines with geographic distance within each season—a critical test for spatial structuring: a significant distance decay would indicate dispersal limitation or spatially structured environmental filtering, whereas weak or absent distance decay would suggest strong mixing and homogenization within the lake.

Second, we aimed to disentangle the ecological processes driving this transition. We asked: (ii) How do the relative contributions of deterministic selection, dispersal, and ecological drift shift from spring to summer across different microbial groups? We addressed this by integrating three complementary lines of evidence: phylogenetic null modeling (βNTI) to detect the signature of homogeneous or variable selection, taxonomic null modeling (the Raup-Crick metric based on Bray-Curtis dissimilarity, RC_bray) to infer the influence of dispersal and drift, and the Sloan neutral community model to estimate the overall fit to neutrality and the inferred migration rate (m). Together, these methods allow us to move beyond a binary deterministic/stochastic classification towards a more nuanced, process-explicit understanding.

Third, we investigated the ecological consequences of this assembly shift. We asked: (iii) How does the spring-summer transition restructure the network of potential interactions within and across microbial groups, and what does this imply for community robustness? We used Sparse Correlations for Compositional data (SparCC) correlation networks to infer co-occurrence patterns and compared their topology (e.g., connectivity, modularity, proportion of positive/negative links) and robustness (simulated species removal) between seasons.

In this study, “cross-domain” refers to an integrative framework that includes two complementary approaches: (i) parallel analysis of each microbial group (e.g. bacterioplankton, cyanobacteria, phytoplankton or zooplankton) to compare their assembly regimes and within-domain network structures (Fig. 1-7, Fig. S3-S10), and (ii) integrated co-occurrence network construction combining multiple groups to reveal potential cross-group interactions (Fig. S11).

**Figure 1.**
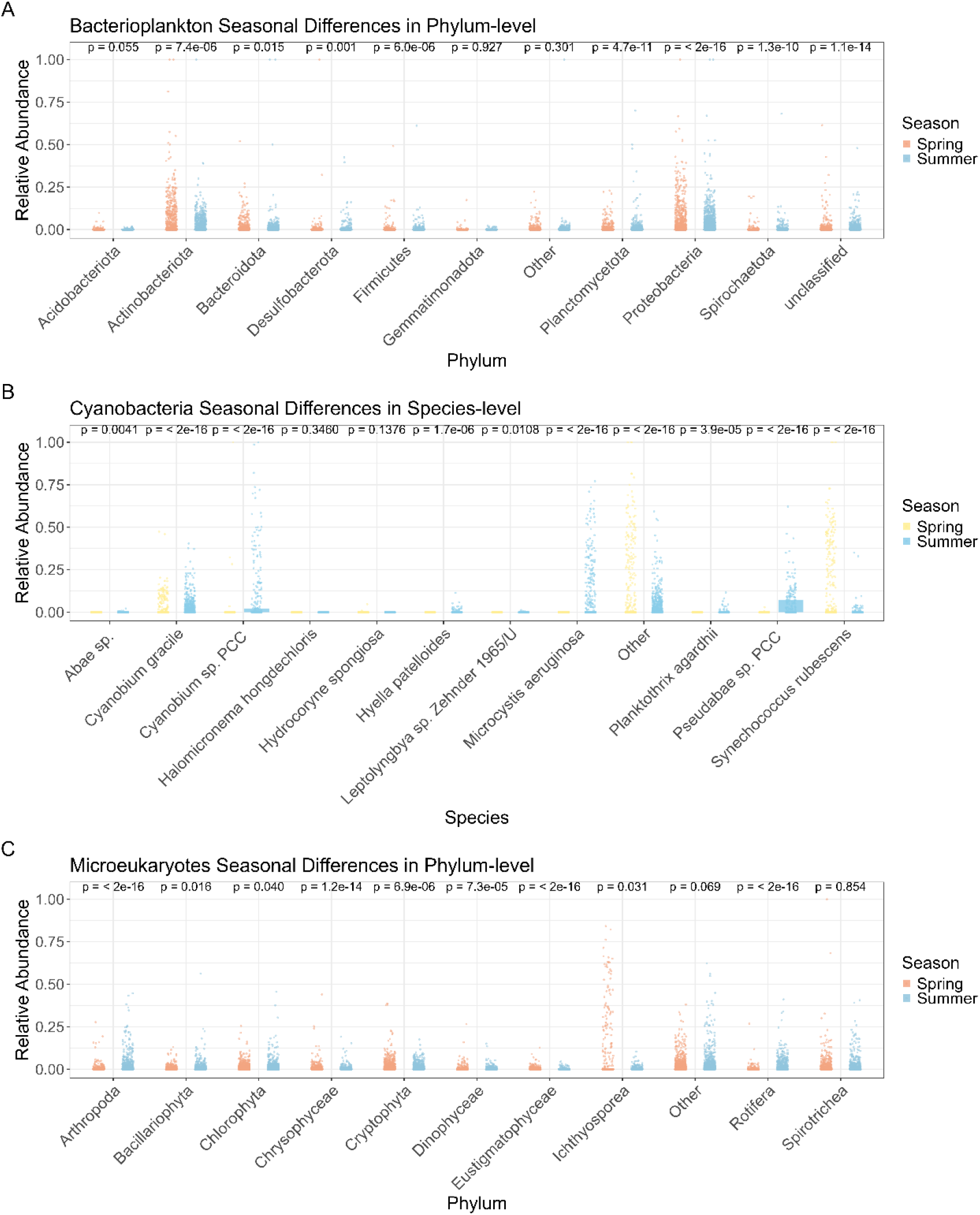
Seasonal differences in relative abundance for (A) bacterioplankton (phylum), (B) cyanobacteria (species), and (C) microeukaryotes (phylum). Shown are the top 10 taxa per group, with others pooled as “Other.” Points show sample-wise relative abundances with overlaid boxplots. p-values from Wilcoxon tests compare seasons.

## Materials and Methods

### Sample collection

Water samples were collected from four phytoplankton-dominated regions of Lake Taihu (North, West, Center, South) during spring (April 1–3) and summer (August 1– 3) 2017 (Fig. S1; Table S1). All samples were from open-lake sites; no riverine inflows were included. The East region, dominated by macrophytes, was excluded. The North and West regions are historically prone to severe algal blooms [41, 42].

We collected 116 composite samples (North 35, West 21, Center 28, South 32). Composite samples combined subsamples from 0.5, 1.0, and 2.0 m to represent the integrated water column. Sites were evenly distributed within each region; only representative locations are shown in Fig. S1. Geographical coordinates were recorded by GPS, and pairwise distances were computed in R using geosphere [44]. Minor variation in sample numbers among regions reflects differences in lake surface area. In this study, we use the term ‘seasonal’ to specifically refer to the contrast between spring and summer—the two critical phases of the bloom transition—rather than implying full annual dynamics.

Samples were pre-filtered through a 200 μm mesh and ∼500 mL was filtered onto 0.2 μm polycarbonate membranes (Millipore, USA). Filters were immediately placed on dry ice, transported to the laboratory within 12 h, and stored at −80 °C until DNA extraction. The broad spatial coverage across 116 sites provided robust biological replication.

Water temperature (WT) was measured in situ using a YSI 6600 V2 sonde (Yellow Springs Instruments, USA). Chemical parameters—including total nitrogen (TN), total phosphorus (TP), ammonium (NH₄⁺-N), and nitrate (NO₃⁻-N)—were analyzed following APHA standard protocols [45].

### DNA extraction and sequencing

We analyzed 232 samples (116 per season) using three amplicon markers: 16S v3– v4, 18S v9, and 23S rRNA (Table S1–S2). DNA was extracted using the OMEGA E.Z.N.A.™ Mag-Bind Soil DNA Kit (OMEGA Bio-Tek, Norcross, GA, USA) with a 50 μL elution volume. PCRs were run in triplicate to reduce stochastic bias, and products were checked by gel electrophoresis.

Each locus was amplified with published primer sets (Table S2) for 16S, 18S, and 23S targets [46–49]. Multiplex PCR conditions, including annealing temperatures and cycling parameters, were optimized by gradient testing and are detailed in Figure S2. Amplicons were purified using SPRI beads (0.8×–1.1× ratio) and pooled in equimolar concentrations. Sequencing was performed on an Illumina MiSeq platform (2 × 300 bp), with a 30% PhiX spike-in to compensate for the low sequence diversity typical of multiplexed amplicon libraries.

### Bioinformatics analyses

Custom reference databases were constructed via in silico PCR using ecoPCR [50] against EMBL [51], allowing ≤3 mismatches. BLAST-based refinement followed a modified CRUX workflow involving retrieval, dereplication, iterative expansion, and taxonomic consolidation [52, 53], which included: (i) in silico PCR retrieval of target-locus sequences; (ii) dereplication; (iii) iterative BLAST-based expansion of the reference set; and (iv) taxonomic consolidation prior to training the Naïve Bayes classifier. Taxonomic assignment of Amplicon Sequence Variant (ASV) was performed using Entrez-QIIME within the QIIME2 framework [54].

Raw reads were demultiplexed, primers trimmed with Cutadapt [55], and sequences were processed with DADA2 [56] for denoising, merging, and chimera removal. Paired-end reads were merged with ≥20 bp overlap, using quality-based truncation to ensure sufficient overlap for all three amplicons. For 18S v9, natural length variation led to some unmergeable pairs, which were retained as high-quality forward reads following the same maxEE and quality-truncation thresholds [48, 57].

Taxonomy was assigned using a Naïve Bayes classifier trained on the custom reference databases. ASVs with confidence scores <0.75 were retained but labeled as “NA” (unclassified) at the corresponding taxonomic rank.

### Statistical analyses

Relative abundances were calculated at the phylum, genus, and species levels, with low-abundance taxa grouped as “Other.” Indicator species were identified using *multipatt()* in the indicspecies package (999 permutations) [58]. Community structure was assessed using non-metric multidimensional scaling (NMDS) based on Bray– Curtis dissimilarities, canonical correspondence analysis (CCA), and permutational multivariate analysis of variance (PERMANOVA) in the vegan package [59–62]. NMDS based on Bray–Curtis dissimilarities was used to visualize overall patterns in community composition. CCA was performed to evaluate relationships between environmental variables and community composition, with the significance of the overall model and canonical axes assessed by 999 permutations. PERMANOVA tested for seasonal and regional differences in community structure. All seasonal subsets were analyzed separately.

To determine whether seasonal community changes are driven by species replacement or by shifts in relative abundances—addressing our first research question—we partitioned Bray-Curtis dissimilarities into turnover (β_bal_) and abundance-gradient (β_gra_) components following Baselga (2013) [17]. β_bal captures compositional replacement, whereas β_gra reflects differences in relative abundances. Partitioning was performed using the betapart package [17, 63, 64]. Because amplicon data are compositional, β_bal and β_gra represent proportional turnover and proportional abundance gradients rather than absolute biomass changes; consequently, β_gra is expected to be small. The ASV table was matched with metadata. Samples with zero total ASV counts (i.e., no sequences recovered) and ASVs with zero counts across all samples were removed. The remaining abundances were then standardized to relative values within each sample using vegan::decoestand (method = ‘total’). Bray–Curtis dissimilarity and its components were computed using *betapart*::beta.pair.abund(). The formulas are:

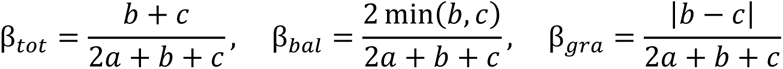

where *a*, *b*, and *c* represent the sums of shared, sample-1-unique, and sample-2-unique abundances, respectively. Differences in β_bal and β_gra among within-season and between-season sample pairs were tested using Wilcoxon rank-sum tests with Benjamini–Hochberg correction (rstatix). Distance–decay patterns (the decline in similarity with increasing geographic distance) were quantified following standard approaches [65, 66]. Bray–Curtis dissimilarities were computed from relative-abundance ASV tables using vegan [67]. and pairwise geographic distances were calculated from sample coordinates using the Haversine formula (geosphere). Mantel tests (999 permutations) were used to assess correlations between community and geographic distance, and ordinary least squares regressions were fitted to estimate slopes, coefficients of determination (R²), and significance.

Environmental predictors were numeric variables standardized to z-scores. Spatial predictors were distance-based Moran’s eigenvector maps (dbMEM) generated from cleaned WGS84 coordinates projected to UTM (sf), using minimum-spanning-tree truncation (igraph) and implemented with adespatial [68–70]. For cross-seasonal variation partitioning, community counts were converted to relative abundances and Hellinger-transformed to generate the response matrix [71]. Within each season (Spring, Summer; n ≥ 6), Redundancy Analysis (RDA; vegan) was used to quantify adjusted R² for pure environment (E|S), pure space (S|E), shared (E∩S), and unexplained fractions; negative fractions were truncated to zero. Significance of pure fractions was evaluated by permutation tests (999 permutations).

To disentangle stochastic and deterministic processes, we followed the two-step framework of Stegen et al. [19, 20], using 999 abundance-based randomizations (vegan [67]). First, we calculated the β-nearest taxon index (βNTI) for each pair of communities within each domain. Pairs with |βNTI| > 2 were interpreted as being governed by homogeneous selection (βNTI < -2) or variable selection (βNTI > +2). For the remaining pairs (|βNTI| < 2), where selection is not the dominant process, we computed the Raup-Crick metric based on Bray-Curtis dissimilarity (RC_bray) to further distinguish the influences of dispersal and drift: RC_bray < -0.95 indicates homogenizing dispersal, RC_bray > +0.95 indicates dispersal limitation, and |RC_bray| < 0.95 reflects undominated processes (primarily drift). Phylogeny-informed turnover was quantified using the β–nearest taxon index (βNTI) for each microbial domain separately. Domain-specific maximum-likelihood phylogenetic trees were constructed from ASV representative sequences for bacterioplankton (16S), cyanobacteria (23S), and microeukaryotes (18S). βNTI was calculated using the abundance-weighted *comdistnt* metric in picante, with null expectations generated by taxa-shuffle randomization (199 permutations). βNTI < −2 indicates homogeneous selection, βNTI > +2 indicates heterogeneous selection, and |βNTI| < 2 indicates that deterministic selection is not the dominant process; in these cases, the relative influences of dispersal and drift were further distinguished using RC_bray. Because 16S, 18S, and 23S target distinct lineages and do not yield a meaningful unified phylogeny, βNTI was computed within domains, consistent with established multi-marker assembly frameworks [26]. Neutral dynamics were assessed using the Sloan neutral model (Sloan et al., 2006) implemented with *neutral.fit()* in the MicEco package, estimating immigration rate (m), metacommunity size (N), and model fit (R²). Taxa were classified based on their occurrence relative to the neutral model prediction: those within the 95% confidence interval were considered neutral (consistent with stochastic processes); those above the upper limit were overrepresented (occurring more frequently than expected, suggesting niche preferences or deterministic selection); those below the lower limit were underrepresented (occurring less frequently than expected, suggesting environmental filtering or competitive exclusion). The model assumes ecological equivalence and equilibrium birth–death–immigration dynamics and is widely applied to quantify neutral contributions in microbial communities [27, 72, 73].

SparCC was used to infer co-occurrence networks for bacterioplankton, cyanobacteria, microeukaryotes and cross-domain assemblages. Because ASV-level SparCC coefficients are small, thresholds (|r| = 0.03–0.07) were chosen based on empirical distributions and network connectivity [74]. Network metrics, including node degree, edge weight, clustering coefficient, transitivity (the global clustering coefficient, measuring the probability that adjacent nodes are connected to each other), path length, and modularity, were calculated using igraph [75], and modules were identified using fast-greedy clustering. For cross-domain network construction, we merged the ASV tables from all three markers (16S, 18S, 23S) by sample, generating a single combined ASV table that included taxa from bacterioplankton, cyanobacteria, and microeukaryotes. This merged table was then used as input for SparCC to infer co-occurrence patterns across domains. Network robustness was assessed by sequential removal of high-degree nodes, calculating the relative size of the largest connected component after each removal, and integrating the robustness curve (area under the curve, AUC). Degree distributions were fitted to power-law models using the poweRlaw package [76].

All analyses were performed in R 4.3.0 [44] using vegan, ggplot2, ggpubr, and related packages. For multivariate analyses (e.g., NMDS, CCA), Bray–Curtis dissimilarities were calculated from relative-abundance matrices. When extreme abundance gradients hindered model convergence, a Hellinger transformation was applied to stabilize variance using *vegan*: decostand (method = “hellinger”).

For eukaryotic microorganisms, we adopted a flexible analytical approach based on the ecological question addressed. For broad-scale community pattern analyses (e.g., NMDS, CCA, β-diversity partitioning), we combined all eukaryotic sequences into a single “microeukaryotes” group to provide an integrated view of total eukaryotic community structure and enable direct comparison with prokaryotic groups. For analyses targeting functional roles and lineage-specific assembly mechanisms (e.g., indicator species analysis, null models, neutral model, co-occurrence networks), we separated eukaryotic sequences into “zooplankton” and “eukaryotic phytoplankton” to capture their distinct ecological responses to bloom conditions.

## Results

### Seasonal and spatial patterns of community composition

ASV-based relative abundance profiles revealed clear seasonal and spatial variation across bacterioplankton, cyanobacteria, and microeukaryotes (Fig. S3). Bacterioplankton were predominantly dominated by Proteobacteria, Actinobacteriota, and Bacteroidota, with modest spatiotemporal variation (Fig. S3A,B). Cyanobacteria showed much stronger seasonality (Fig. S3C,D): spring communities were dominated by *Synechococcus rubescens*, accompanied by high abundances of *Cyanobium gracile*, whereas summer communities were led by *Microcystis aeruginosa*, with *Pseudanabaena* spp. strongly increased—especially in the West—while *C. gracile* and other *Cyanobium* spp. remained present at lower relative abundances. Microeukaryotes showed the greatest spatial heterogeneity (Fig. S3E,F), with spring assemblages dominated by Cryptophyta, Chlorophyta, and regionally enriched Bacillariophyta. In summer, Cryptophyta remained abundant in the Center/West, Chlorophyta increased in the South, Bacillariophyta expanded in the North, and Dinophyceae increased in the West. Unassigned taxa were kept at class level, and only phytoplankton and zooplankton were included.

Seasonal shifts in dominant groups (Fig. 1) showed significantly higher summer abundances of Proteobacteria, Actinobacteriota, and Bacteroidota in bacterioplankton (p < 0.01; Fig. 1A). Cyanobacteria exhibited the strongest turnover, shifting from *S. rubescens*, *C. gracile*, and *Cyanobium* spp. in spring to *M. aeruginosa* and *Pseudanabaena* spp. in summer (p < 0.05; Fig. 1B). Microeukaryotic phyla varied more subtly, with Ichthyosporea consistently enriched in spring (Fig. 1C), matching spatial patterns in Fig. S3E–F. Detailed taxonomic compositions at phylum, genus, and species levels for both seasons are provided in Tables S3 and S4.

Indicator species analysis (Fig. S4; p < 0.01) further supported seasonal differentiation. Spring bacterioplankton indicators included *Bacteriovorax* sp. and *Saccharimonas* sp., whereas summer featured *Enterobacteriaceae* bacterium, *Aurantimonadaceae* bacterium, and *Candidatus Aquiluna* sp. (Fig. S4A). Cyanobacterial indicators shifted from *S. rubescens* in spring to *M. aeruginosa* and *Cyanobium* spp. in summer (Fig. S4B). Zooplankton indicators comprised *Sinocalanus tenellus*, *Synchaeta pectita*, and *Myxophyllum magnum* in spring, versus *Eurycercus lamellatus*, *Hexarthra intermedia*, and *Bosmina longirostris* in summer (Fig. S4C). Eukaryotic phytoplankton showed parallel changes: spring indicators such as *Paraphysomonas uniformis*, *Aulacoseira subarctica*, and *Heterosigma rufescens* typified cooler seasons, whereas summer indicators (*Aulacoseira granulata*, *Mychonastes* sp., *Phacotus lenticularis*) reflected warmer, more eutrophic conditions (Fig. S4D).

### Seasonal ordination and β-diversity partitioning

NMDS ordinations showed clear seasonal separation of bacterioplankton, cyanobacteria, and microeukaryotes (Fig. 2A–C; stress = 0.131, 0.092, 0.115). PERMANOVA revealed strong seasonal shifts in all groups, while within-season dissimilarities remained high (median > 0.80), indicating substantial heterogeneity (Table S5). Effects of sampling region (North, West, Center, South) were small (R² = 0.015–0.039) and mostly non-significant or weak, whereas a modest but significant Season × Region interaction (R² ≈ 0.018; p = 0.008) indicated slight shifts in spatial structuring between spring and summer. Region effects were weak in spring (p = 0.062– 0.218) but became marginally significant for some groups in summer (p = 0.016–0.097), although these remained minor compared with the dominant seasonal signal. CCA identified water temperature as the dominant seasonal driver, with NO₃⁻ additionally influencing cyanobacteria and microeukaryotes (Fig. 2G–L). Spatial CCA within seasons (Fig. S5) explained < 2.5% variation with no significant predictors, indicating minimal measured environmental control at within-season scales.

**Figure 2.**
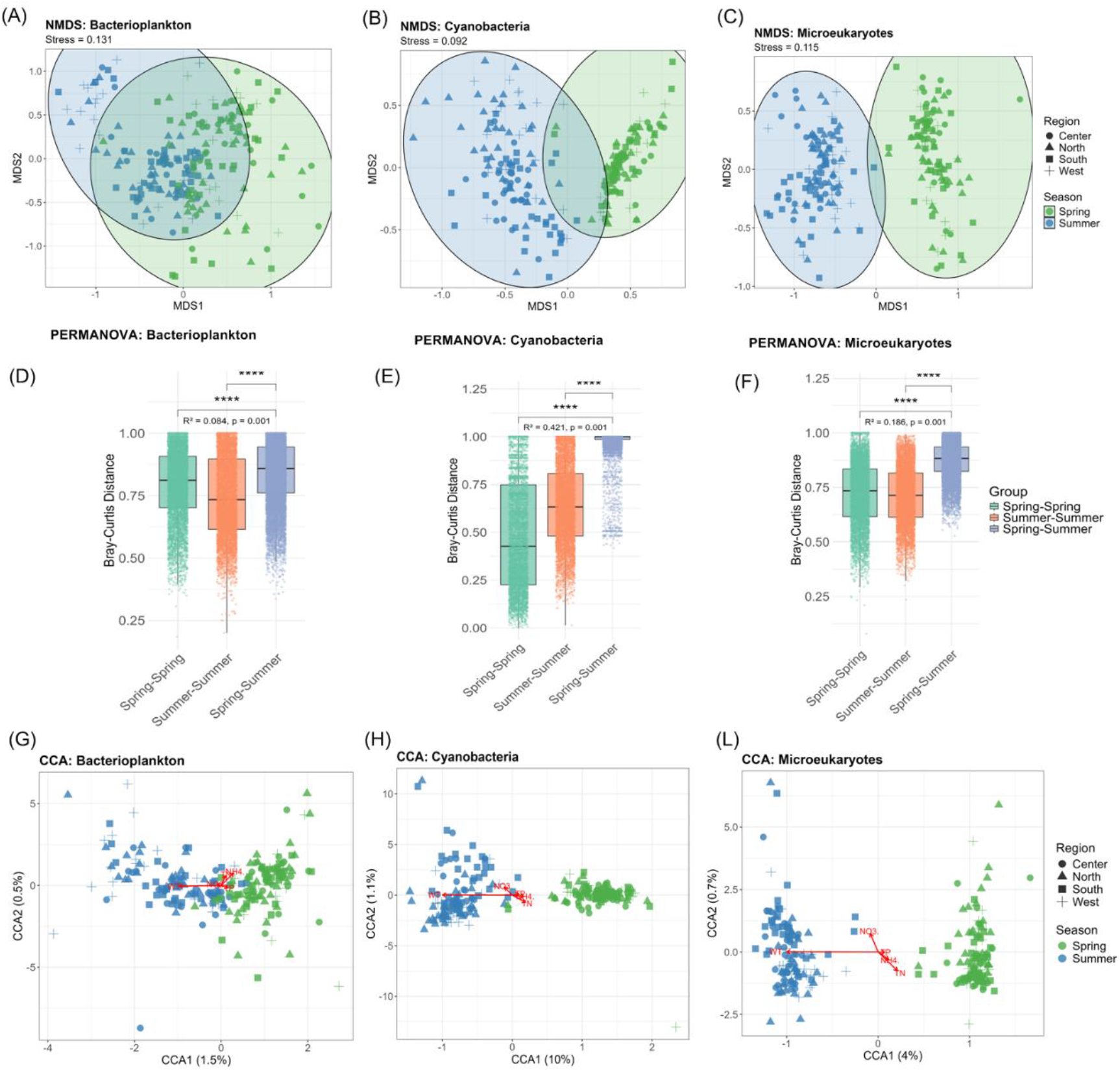
Community patterns of bacterioplankton, cyanobacteria, and microeukaryotes between seasons. (A–C) NMDS ordinations (Bray–Curtis), colored by season and shaped by region; ellipses indicate dispersion (stress shown). (D–F) CCA ordinations constrained by environmental variables. (G–H) Pairwise Bray–Curtis dissimilarities with Wilcoxon tests; PERMANOVA R² and p-values shown.

β-diversity partitioning (Fig. 3) showed Bray–Curtis dissimilarity was almost entirely due to turnover (β_bal), with negligible abundance-gradient effects (β_gra). Within-season spatial contrasts were significant for bacterioplankton in both seasons (p < 0.001), but only in summer for cyanobacteria and microeukaryotes. For all groups, cross-season turnover greatly exceeded within-season turnover (p < 0.001), demonstrating that seasonal transitions intensified species replacement beyond spatial effects alone.

**Figure 3.**
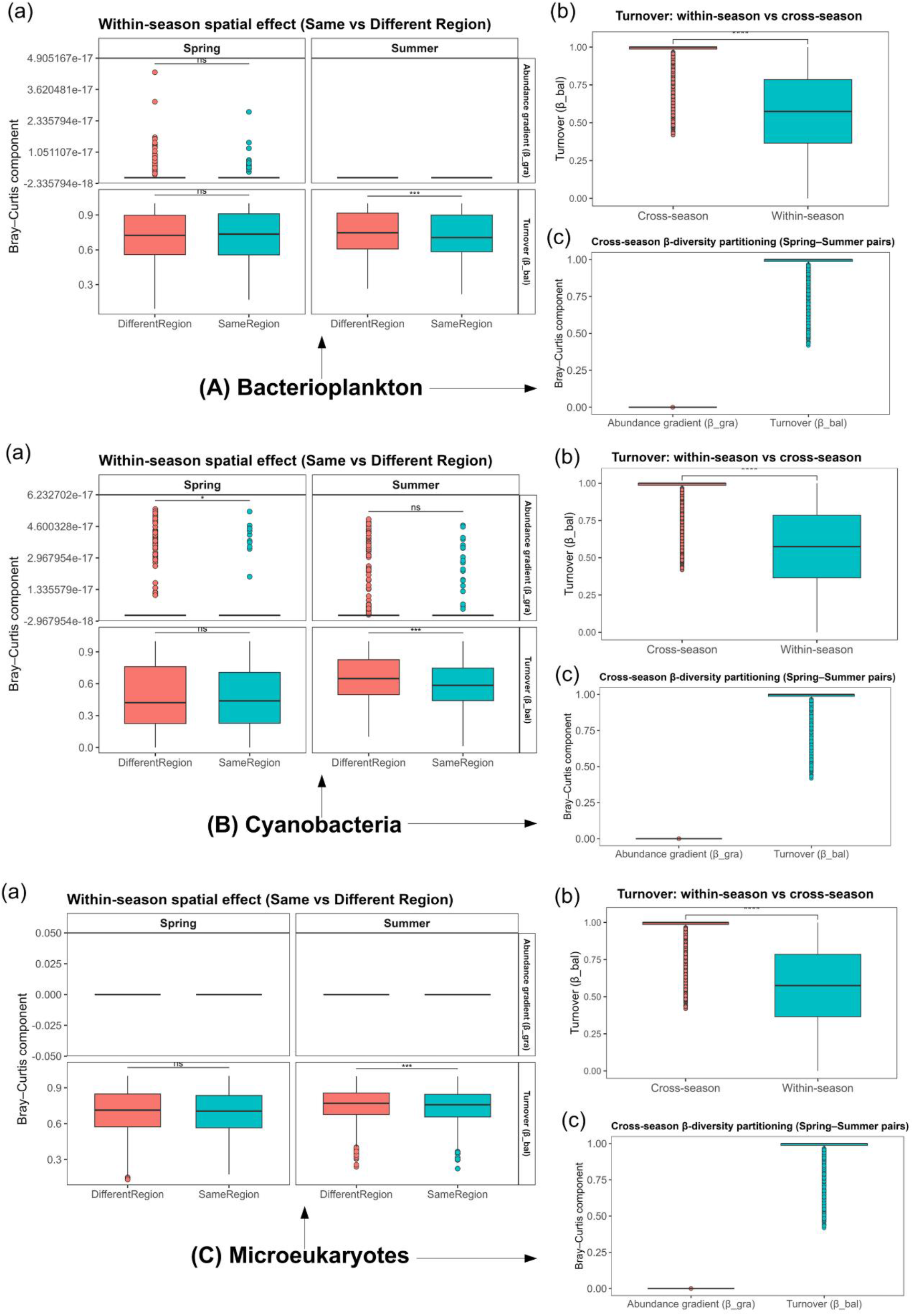
β-diversity partitioning for bacterioplankton, cyanobacteria, and microeukaryotes. For each group: (a) within-season spatial contrasts by region; (b) within-vs. cross-season turnover; (c) β-diversity decomposition showing that seasonal differences are driven mainly by turnover. *p < 0.001; ns, not significant.

### Variation partitioning and distance–decay

The variation partitioning analysis separated the explained variation into pure environmental (E|S), pure spatial (S|E), shared (E∩S), and unexplained fractions. Variation partitioning (Fig. 4A–C) showed pure environmental and spatial factors explained very little within-season variation (<4%). Bacterioplankton showed E|S/S|E contributions of ∼1.0%/3.4% in spring and ∼1.9%/2.2% in summer. Cyanobacteria increased from 0%/6.4% in spring to 3.1%/12.3% in summer, whereas microeukaryotes remained low (spring: 0.9%/3.0%; summer: 1.0%/3.3%). Full results (Fig. S6) confirmed the dominance of unexplained variation (∼95–97% in spring; ∼93–96% in summer; cyanobacteria: 94% spring, 88% summer), with shared E∩S consistently minimal.

**Figure 4.**
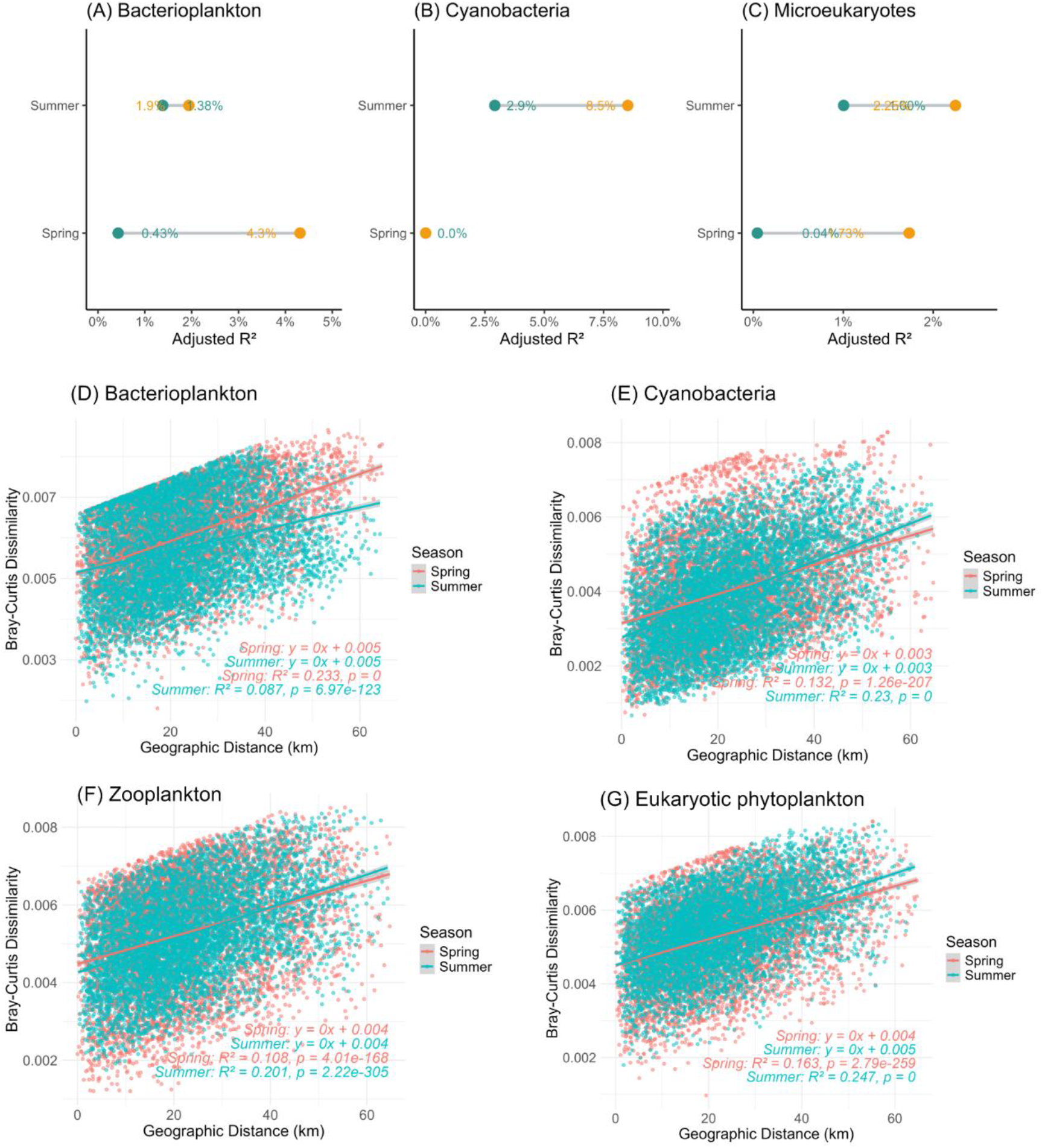
Environmental vs. spatial contributions and distance–decay. (A–C) Variation partitioning from partial RDA comparing pure environmental and pure spatial/dbMEM fractions. (D–G) Distance–decay relationships (Bray–Curtis vs. geographic distance) with seasonal regressions and R² and p-values. These analyses highlight seasonal shifts in environmental filtering and spatial structuring.

Distance–decay analyses (Fig. 4D–G) showed significant positive relationships between community dissimilarity and geographic distance in both seasons (p < 0.001). Bacterioplankton showed stronger decay in spring (R² = 0.233) than summer (R² = 0.087), whereas cyanobacteria, zooplankton, and eukaryotic phytoplankton exhibited stronger decay in summer (cyanobacteria: 0.132→0.230; zooplankton: 0.108→0.201; eukaryotic phytoplankton: 0.163→0.247). All slopes were shallow, reflecting gradual spatial differentiation.

### Null- and neutral-model

βNTI patterns (Fig. 5A–C) showed domain-specific deterministic signatures. Bacterioplankton values largely clustered within |2|, indicating near-neutral dynamics. Cyanobacteria exhibited strongly negative βNTI values in spring (median < -2), indicating dominance of homogeneous selection, and shifted toward less negative values in summer while remaining selection-dominated. Microeukaryotes showed negative βNTI in both seasons, with a moderate summer shift toward neutrality. RC_bray results (Fig. 5D–G) paralleled these trends: bacterioplankton centered near zero; cyanobacteria and zooplankton displayed many values < −0.95; eukaryotic phytoplankton centered near zero with a distinct negative tail.

**Figure 5.**
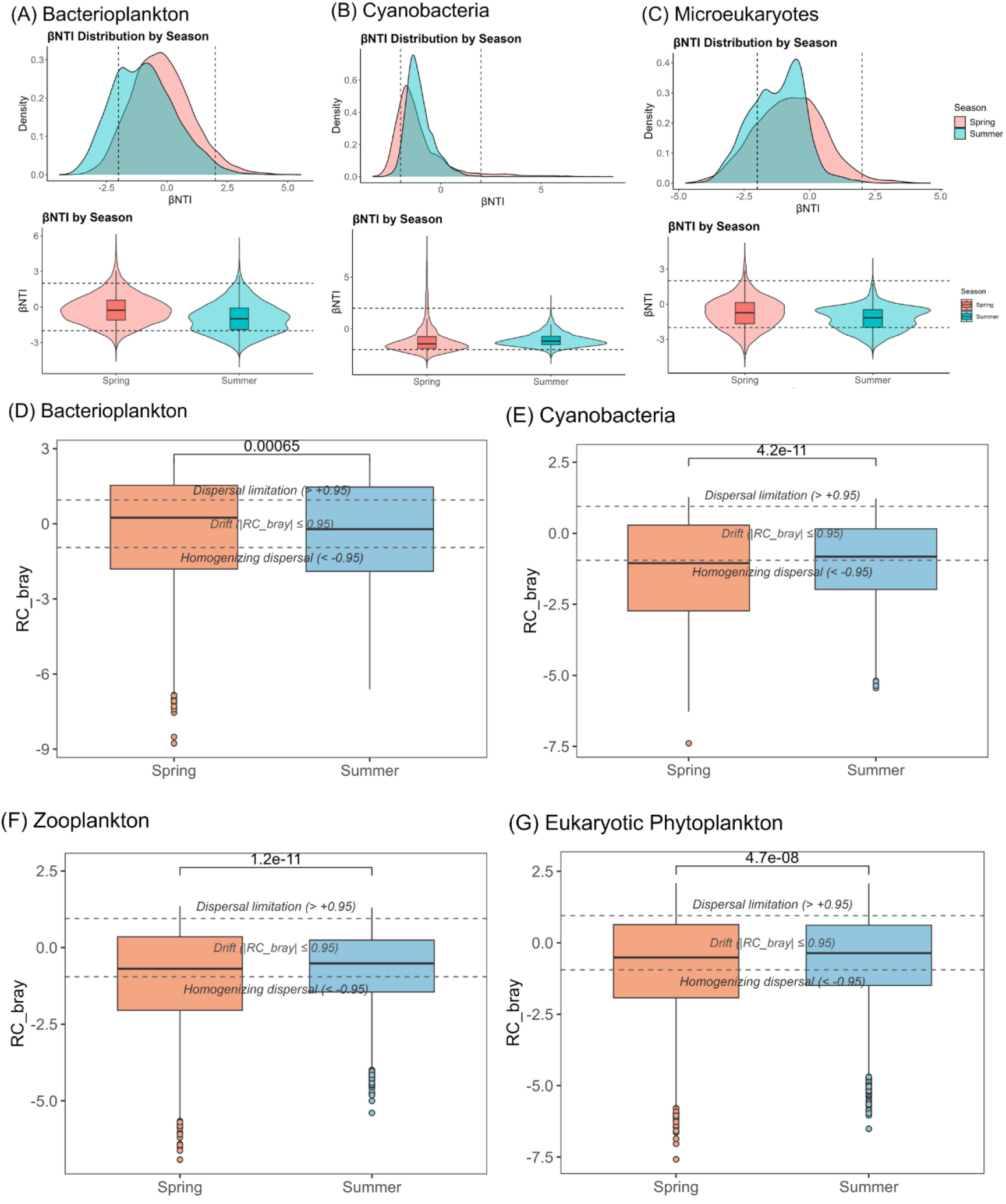
Null-model analyses of bacterioplankton, cyanobacteria and microeukaryotes assembly. (A–C) βNTI distributions from 199 phylogenetic null permutations; dashed lines at ±2 mark deterministic thresholds. (D–G) RC_bray from 199 abundance-shuffled null communities; dashed lines at ±0.95 indicate dispersal limits. Together these metrics quantify seasonal differences in deterministic vs. stochastic processes.

Neutral-model fits (Fig. S7) showed high bacterioplankton neutrality (R² > 0.87) with higher Nm in summer. Cyanobacteria showed weak neutral fits in both seasons, with even more deviation from neutrality in summer. Eukaryotic phytoplankton showed moderate fits (spring R² = 0.761; summer R² = 0.738), with neutrality increasing from 84.7% to 97.4% and Nm decreasing from 14.105 to 10.138.

### Network-based community interactions

Network topology and robustness showed clear spring–summer reorganization across within-domain and cross-domain networks (Fig. 6–7). Bacterioplankton networks expanded modestly with minimal changes in topology or interaction polarity. Cyanobacterial networks underwent the strongest restructuring: node and edge counts increased substantially, associations shifted from predominantly negative in spring to more positive in summer, and robustness increased markedly. Microeukaryotic networks increased modestly in size while remaining dominated by negative associations. Cross-domain networks expanded substantially, with a pronounced shift toward positive associations and increased robustness (Fig. 7D).

**Figure 6.**
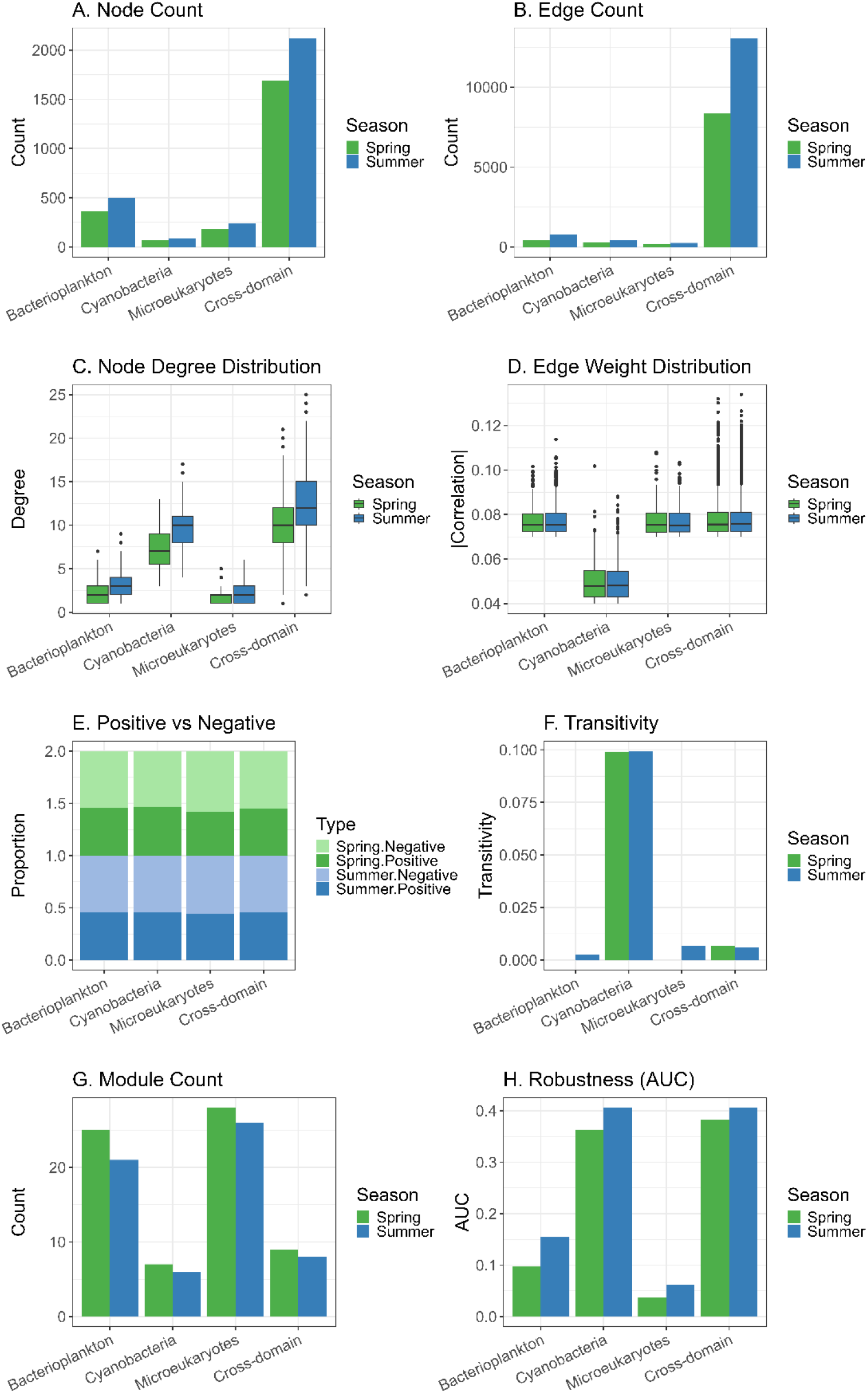
Seasonal differences in SparCC network structure for bacterioplankton, cyanobacteria, microeukaryotes and cross-domains. (A–B) Node and edge counts. (C– D) Degree and edge-weight distributions. (E) Positive vs. negative correlations. (F) Global transitivity. (G) Module numbers. (H) Robustness (AUC) under targeted node removal. Metrics highlight seasonal changes in network size, connectivity, and stability.

**Figure 7.**
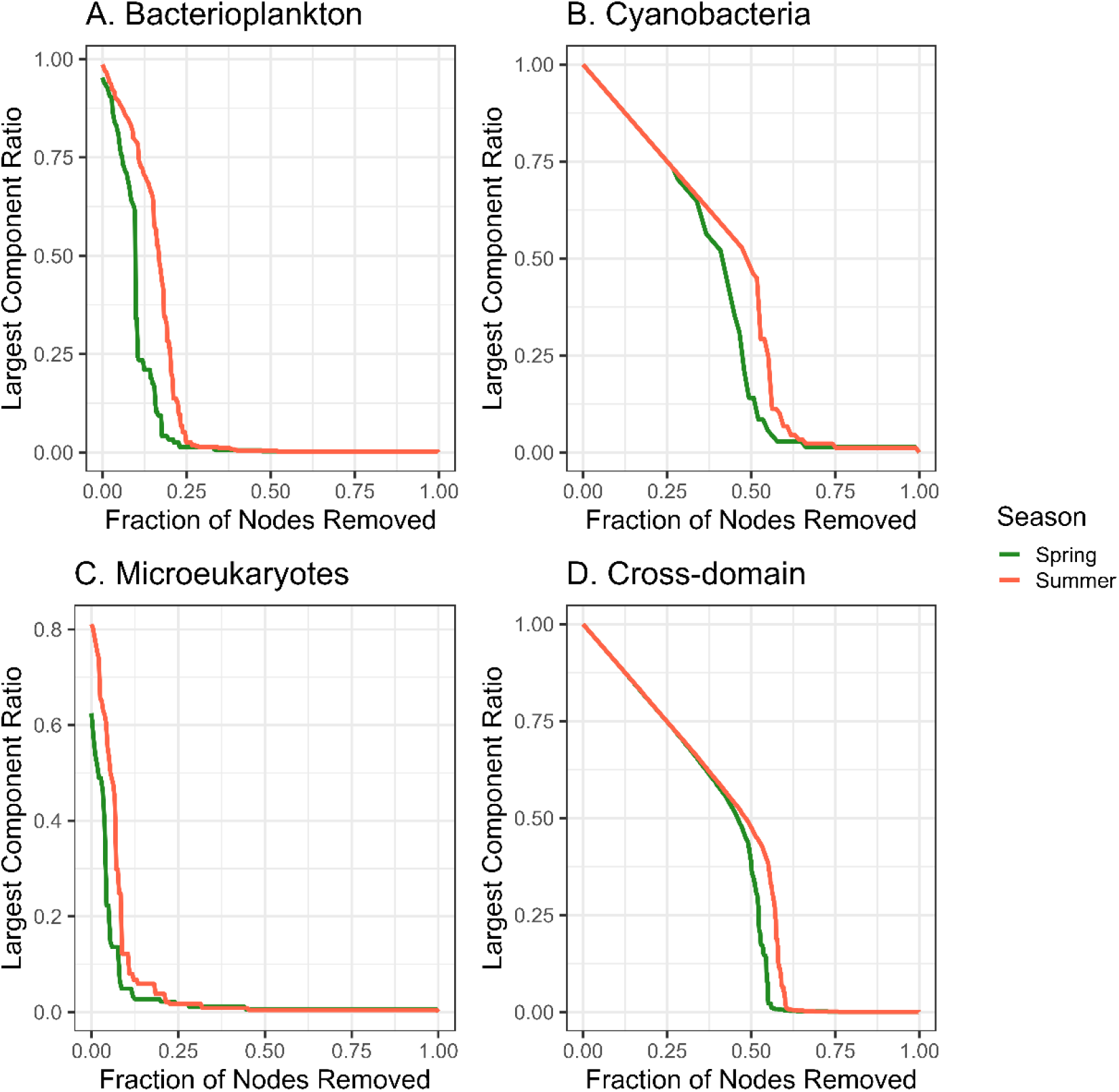
Network robustness for bacterioplankton (A), cyanobacteria (B), microeukaryotes (C), and cross-domain networks (D). Curves show the retained largest-component size as high-degree nodes are removed; steeper slopes indicate lower robustness.

Domain-specific networks (Fig. S8–S10) supported the seasonal patterns: bacterioplankton networks showed stronger inter-phyla connectivity; cyanobacterial networks formed more cohesive summer modules centered on Microcystis; microeukaryotic networks incorporated broader taxonomic representation (Ascomycota, Rotifera). Note that for cyanobacteria, the 23S marker recovered relatively few cyanobacterial ASVs, making a robust cyanobacteria-only network unfeasible. We therefore constructed a cross-kingdom network (Fig. S9) that included all taxa (bacteria, cyanobacteria, and microeukaryotes) to capture potential interactions between cyanobacteria and other microbial groups, leveraging the 23S marker’s ability to simultaneously amplify cyanobacteria and eukaryotic plastids. Cross-domain networks (Fig. S11) shifted from more negative to more positive associations, with intensified cyanobacteria–eukaryote links (e.g., Synechococcales/Nostocales with Actinobacteriota, Proteobacteria, Chlorophyta, Spirotrichea).

Species-level networks (Fig. S12) highlighted contrasting bloom-stage roles: spring *S. rubescens* associated negatively with *Aphanomyces invadans* and positively with *Ergasilus scalaris*; summer *M. aeruginosa* showed negative associations with rotifers and amoebozoans (*Vannella simplex*, *Diplophrys mutabilis*, *Hexarthra intermedia*) and positive links with *Synchaeta pectita* and *Flavobacterium* spp., suggesting potential ecological interactions that warrant experimental validation.

## Discussion

In this study, we conducted a large-scale, multi-marker eDNA assessment of synchronous seasonal dynamics across prokaryotic and eukaryotic microplankton in Lake Taihu during 2017—one of the most extensive bloom years in recent decades. By integrating multi-marker diversity, β-diversity partitioning, distance–decay relationships, null and neutral model analyses, and network analyses, we resolved how filtering, dispersal, and stochasticity shaped seasonal reassembly and restructuring. Extending earlier single-domain studies in Taihu and other eutrophic lakes [10, 11, 28, 77], our cross-domain framework reveals contrasting assembly regimes and seasonal reorganization of interaction architecture [40–42]. These findings provide new insights into how seasonal environmental filtering and bloom-driven resource reconfiguration jointly govern cross-domain assembly and interaction restructuring in eutrophic lake ecosystems.

### Seasonal microbial composition shifts

Bacterioplankton were dominated by Proteobacteria, Actinobacteriota, and Bacteroidota, but indicator taxa shifted from *Bacteriovorax* and *Saccharimonas* in spring to members of Enterobacteriaceae bacterium, Aurantimonadaceae bacterium, and *Candidatus Aquiluna* in summer, reflecting selection under warmer, nutrient-rich conditions [4, 78]. Cyanobacteria shifted from *Synechococcus rubescens* and *Cyanobium gracile* in spring to *Microcystis aeruginosa* in summer, especially in central and southern regions. Unlike Chaohu—where picocyanobacteria collapse—*C. gracile* and *Cyanobium* spp. remained abundant in Taihu, especially in the North and West, consistent with tolerance to fluctuating light/nutrients and particle-associated niches [10, 17].

Turbulence-driven trait selection may affect *Microcystis* photophysiology [12]. Whereas *Cyanobium* declines sharply in Chaohu [79], Taihu retained abundant *Cyanobium* and showed strong summer increases of filamentous *Pseudanabaena*, especially in the West [80]. This taxon, likely *P. galeata* (Table S4), is known to thrive under stratified and turbid conditions as a low-light specialist, contributing to its summer proliferation. The high-resolution multi-marker eDNA approach and fine-scale resource patchiness may further facilitate its coexistence with other cyanobacteria [31, 32]. Frequent *Pseudanabaena*–Microcystis co-occurrence likely reflects shared mucilage rather than direct interactions [6–8, 36, 81], consistent with *P. mucicola* (a species known to live embedded within *Microcystis* colonies [81]), while free-living *Pseudanabaena* filaments frequently co-occur with *Microcystis.* Spatial heterogeneity in nutrient availability may further shape phytoplankton community structure. Summer phytoplankton indicators (*Aulacoseira granulata*, *Mychonastes* sp., *Phacotus lenticularis*) were positively associated with NH₄⁺ (Fig. S4D), while within-season CCA explained little variation (Fig. S5), suggesting that reduced nitrogen forms may contribute to regional differences in community composition, particularly in areas with higher nutrient input [41, 43]. Eukaryotic phytoplankton also showed pronounced seasonal turnover: cooler-water spring taxa (*Paraphysomonas uniformis*, *Aulacoseira subarctica*, *Heterosigma rufescens*) were replaced by bloom-adapted species (*A. granulata*, *Mychonastes* spp., *Phacotus lenticularis*) [42, 82], mirroring cyanobacterial transitions under elevated temperature, stratification, and nutrient enrichment [83]. Similar patterns occur in peri-Alpine and Donghu lakes [10, 11]. Synchronous domain-level changes indicate trophic coupling: cyanobacteria and phototrophs share thermal niches, efficient phosphorus uptake, small size, buoyancy traits, grazer resistance [82, 84–86], and phenological synchrony with temperature/irradiance cues [84]. Zooplankton tracked bloom development, as spring taxa were replaced by cyanotoxin-tolerant rotifers (*E. lamellatus*, *H. intermedia*, *B. longirostris*), with *B. longirostris* increasing during toxic episodes [87–89].

Temperature and nutrient enrichment therefore acted as primary filters structuring seasonal turnover [3, 5], consistent with satellite and machine learning (ML) records from Lake Erie showing seasonal–interannual modulation of bloom extent [7, 8]. Our results reveal coordinated trophic shifts, persistent *Cyanobium*, cyanobacterial– phototroph replacements, and strong rotifer responses—demonstrating restructuring during the spring–summer transition. These findings extend insights from Lake Mendota and Lake Erie by embedding bacterial and zooplankton turnover into a cross-domain seasonal framework.

### Community assembly driven by season-specific interplay of deterministic and stochastic Processes

Our integrative framework (NMDS, CCA, β-diversity partitioning, RC_bray, neutral models) demonstrated coherent, taxon-specific shifts in assembly mechanisms across the spring–summer transition, consistent with niche–neutral continuum theory [14]. Unlike earlier single-domain or long-term studies [10, 11], the cross-domain approach captured synchronous restructuring of bacteria, cyanobacteria, and microeukaryotes.

Seasonal clustering (p < 0.001) and turnover-dominated β-diversity showed that temporal forcing far exceeded spatial heterogeneity [41, 43]. Between-season turnover consistently exceeded within-season contrasts, paralleling the 2017 bloom expansion [100] and other large-lake regime shifts [3, 41, 43, 84, 90, 91]. Evidence from epiphytic and sediment studies [77] similarly highlights the strong imprint of short seasonal windows. Temperature and nutrients were dominant drivers—especially for cyanobacteria—while mixing modulated bloom trajectories [12]. Within-season CCA explained < 2.5% of variation, and variation partitioning indicated extremely high unexplained fractions (≈95–97% in spring; ≈88–96% in summer), confirming weak coupling with measured environmental gradients.

Distance–decay slopes were shallow yet significant across groups, with minimal pure spatial effects for bacterioplankton and microeukaryotes but stronger spatial structuring for cyanobacteria, consistent with dispersal limitation during bloom expansion. This spatial pattern likely reflects the influence of nutrient loading from the drainage basin, as the northwestern region (Zhushan Bay, Meiliang Bay) receives high nutrient inputs from inflowing rivers, creating favorable conditions for bloom initiation and leading to persistent spatial heterogeneity in cyanobacterial biomass [41, 43]. The significant distance-decay relationships indicate the presence of spatial structure in community composition, although the modest R² values (0.087–0.247) are consistent with the small pure spatial fractions (S|E) in the variation partitioning analysis, suggesting that spatial structure, while detectable, explains only a limited portion of community variation after accounting for environmental factors. Such weak but persistent signals typify shallow lakes where mixing homogenizes water masses but bloom development imposes transient spatial structure. Micro-patchiness of labile substrates, particle-associated “lake snow” [31, 32], and basin-scale disturbances [84] likely contribute to this interplay. βNTI patterns aligned with these trends. Cyanobacteria experienced strong homogeneous selection in spring (βNTI < −2), indicating that environmental filtering dominated community assembly during the early bloom period. The shift toward near-neutral βNTI values in summer reflects a relaxation of selection pressure, likely driven by enhanced dispersal due to wind-induced mixing during the bloom stage—a pattern supported by the concurrent increase in RC_bray values toward neutrality. Microeukaryotes showed a similar relaxation of selection, whereas bacterioplankton remained consistently near neutral, suggesting that stochastic processes (dispersal and drift) play a more persistent role in structuring bacterial communities regardless of season. RC_bray corroborated these trends: cyanobacteria and zooplankton exhibited many values < −0.95 in spring, indicating strong homogenizing dispersal during this period, consistent with the well-mixed conditions typical of temperate lakes in early spring, while bacterioplankton centered near zero. Neutral-model fits showed high neutrality for bacterioplankton (R² > 0.87) with higher Nm in summer; cyanobacteria showed weaker fits and more outliers; and eukaryotic phytoplankton exhibited increased neutrality (84.7% → 97.4%) but reduced Nm, suggesting that homogenization in this group was more drift-than dispersal-driven [73, 92].

Overall, Cyanobacteria and microeukaryotes were under strong homogeneous selection in spring but shifted toward drift-like or near-neutral assembly in summer, whereas bacterioplankton stayed near neutral. Thus, deterministic filtering from warming and nutrients drives major seasonal restructuring, while within-season variation reflects stochasticity shaped by lineage traits and dispersal capacity. These patterns align with theoretical frameworks that emphasize the hierarchical interplay of ecological processes [14, 73, 93]: deterministic environmental filtering (selection) drives broad-scale seasonal reorganization, while within-season variation is shaped by stochastic processes (drift and dispersal) whose relative importance is modulated by lineage traits and dispersal capacity. This hierarchy—with selection operating at larger temporal scales and drift/dispersal at finer scales—is consistent with predictions from niche theory.

### Network-based community interactions reveal seasonal restructuring

By integrating multi-marker datasets, we constructed season-specific co-occurrence networks across bacteria, cyanobacteria, phytoplankton, and zooplankton, extending earlier single-domain lake studies that only examined seasonal restructuring within prokaryotes or eukaryotes [10, 14, 15, 17, 28]. Seasonality dominated network reorganization in Taihu, consistent with long-term Mendota patterns [94]; both within-domain and cross-domain networks became larger and more connected in summer, reflecting enhanced resource fluxes and bloom-driven selection [3, 93], though restructuring varied across groups.

Bacterioplankton networks expanded moderately with largely stable degree distributions and positive–negative proportions. Microeukaryotic networks grew slightly, remained dominated by negative associations, and showed only small summer increases in transitivity and robustness. Cyanobacteria underwent the strongest restructuring: substantial increases in nodes, edges, degree, and positive associations, higher robustness, and cohesive summer modules around *Microcystis*, *Trachydiscus*, and *Lepocinclis* (Fig. S8–S10). Cross-domain networks showed the clearest shifts in interaction polarity, transitioning from predominantly negative associations in spring to predominantly positive associations in summer (Fig. 6E). Although similar in size, summer cross-domain networks showed longer path lengths, lower transitivity, fewer negative edges, and more positive associations, indicating reduced antagonism and stronger interdomain coupling [14, 17, 84]. Robustness rose markedly (higher AUC; Fig. 7D). Cyanobacteria–eukaryote links intensified, particularly between Synechococcales/Nostocales and Actinobacteriota, Proteobacteria, Chlorophyta, and Spirotrichea (Fig. S11).

Species-level patterns highlighted contrasting roles of *Synechococcus rubescens* and *Microcystis aeruginosa* (Fig. S12). In spring, the *S. rubescens*-centered network showed negative associations with *Aphanomyces invadans* and Bacteroidetes and positive associations with *Ergasilus scalaris* and a decomposer bacterium. In summer, the *M. aeruginosa*-centered network showed negative associations with *Vannella simplex*, *Diplophrys mutabilis*, *Hexarthra intermedia*, and *Sphaeroplea annuli*, and positive interactions with *Synchaeta pectita*. Strong summer partners—including *Actinobacterium*, *Flavobacterium* spp., *Vorticella microstoma*, *H. intermedia*, and *Paraphelidium tribonemae*—suggest mutualistic, predatory, or parasitic interactions consistent with toxin-mediated zooplankton suppression [3, 22, 95, 96]. Actinobacteria linked broadly with multiple cyanobacterial orders rather than specifically with *Microcystis* (Fig. 6–7). Other bacterial groups, including *Flavobacterium*, *Polynucleobacter*, and *Bacteriovorax* (Table S4), also exhibited strong associations with cyanobacterial taxa, consistent with their known roles as cyanobacteria-associated bacteria in eutrophic lakes.

Overall, the spring–summer transition intensified intra-domain interactions and rebalanced cross-domain connectivity, producing more connected and robust summer networks. This shift likely reflects increased metabolic interdependence and resource exchange under bloom conditions, reinforcing cross-domain coupling and enhancing system-level stability. Shifts in polarity, module cohesion, and hub taxa reflect adaptive reconfiguration that supports functional resilience under bloom-driven abiotic–biotic feedbacks [21, 22, 97]. Nonetheless, compositional data constrain ecological interpretation [3, 21, 74, 98–100], and two-season sampling limits temporal inference, highlighting the need for high-frequency and experimental validation. Importantly, the co-occurrence networks presented here are based on correlation patterns derived from relative abundance data and should be interpreted as hypotheses about potential associations rather than confirmed biological interactions. Such analyses cannot distinguish between direct ecological interactions (e.g., mutualism, predation, competition) and indirect associations arising from shared environmental preferences or common responses to environmental drivers [21, 22]. Experimental validation— such as co-culture experiments or targeted manipulation—is therefore essential to confirm the nature of these inferred associations.

## Implications for lake management and future research

Our results suggest that the effectiveness of management interventions may depend on seasonal assembly regimes. Spring communities, dominated by deterministic selection, may respond more predictably to nutrient reduction and early-season biomanipulation (e.g., enhancing grazer populations). In contrast, summer communities exhibit stronger stochasticity and internal nutrient recycling, implying that disruption of key microbial interactions—such as *Microcystis*-associated consortia— may be more effective than nutrient control alone during bloom peaks (Fig. S13). Network robustness metrics (e.g., modularity, connectivity) and neutrality diagnostics could serve as early-warning indicators of regime shifts, informing adaptive management strategies. These findings highlight the importance of considering cross-domain interactions and seasonally shifting assembly mechanisms when predicting ecosystem responses to eutrophication and climate change. Future research integrating metatranscriptomics and experimental manipulations is needed to validate the ecological mechanisms inferred from our network and assembly analyses.

## Conclusion

This multi-marker eDNA study provides an integrated, cross-domain assessment of microbial community assembly during the spring–summer bloom transition in Lake Taihu. Despite substantial spatial heterogeneity, seasonal turnover dominated across bacterioplankton, cyanobacteria, phytoplankton, and zooplankton. Assembly mechanisms shifted systematically: strong homogeneous selection in spring, particularly for cyanobacteria and microeukaryotes, transitioned toward more drift- and neutrality-like dynamics in summer, aided by enhanced dispersal under bloom-driven mixing. Co-occurrence networks displayed pronounced seasonal restructuring, with summer communities forming more cohesive, positively coupled, and markedly more robust interaction structures. Together, these findings suggest that bloom progression restructures aquatic microbiomes through a seasonal rebalance of selection, drift, and dispersal, coupled with adaptive reinforcement of cross-domain interactions. Such coordinated seasonal dynamics may contribute to functional stability and robustness in eutrophic lake ecosystems under bloom pressure.

## Availability of data

The NGS sequence data of all samples produced in this study are deposited in NCBI. The SRA accession for the NGS sequences is PRJNA909005 (released after acceptance of this manuscript). All custom bioinformatics and R scripts are available at: https://github.com/zousm912zou/Taihu_Microbiome_eDNA_Assembly.

## Authors Contributions

Shanmei Zou conceived and designed the study, performed data analyses, and wrote the manuscript. Shanmei Zou, Lydia Smith, and Xuemin Wu conducted the experiments. Xinke Yu and Tiantian Sun collected the samples. Beixin Wang provided methodological guidance. Tyler Linderoth assisted with experiments and commented on the manuscript draft. Sven Katzenmeier assisted with data analysis and commented on the draft. Thorsten Stoeck contributed to study design and conceptual development and commented on the draft.

## Acknowledgement

This work was supported by the National Natural Science Foundation of China (31600294). Bioinformatics support was provided by the Bioinformatics Center of Nanjing Agricultural University. We thank the Evolutionary Genetics Lab at the University of California, Berkeley, for assistance with NGS library preparation. We also thank Meng Wang and Yuxiao Hua for technical help and the Laboratory of Ecosystem Ecology at Nanjing Agricultural University for experimental platform support.

---

**Figure S1.**
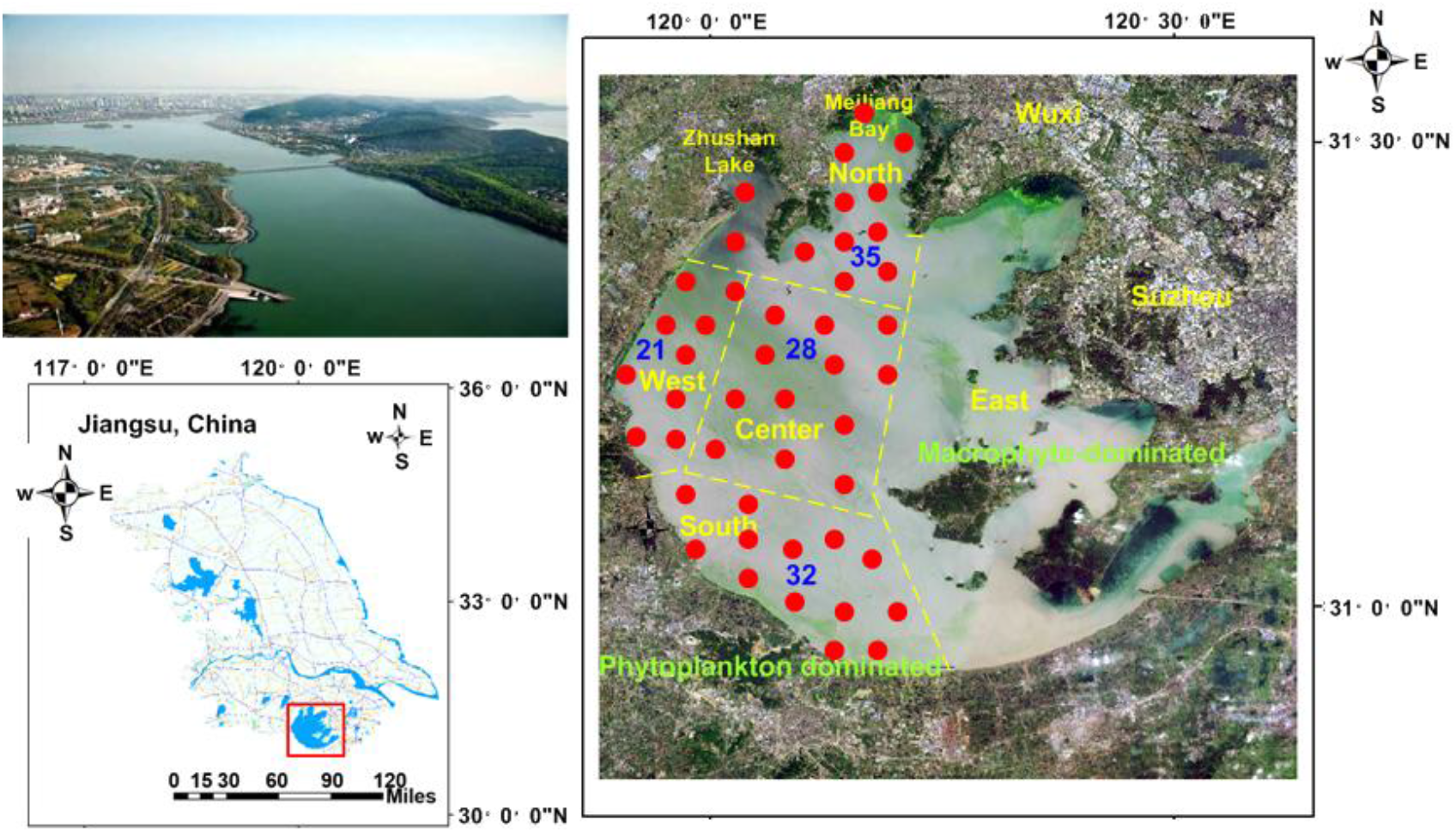
Sampling map of Lake Taihu showing representative sites from four phytoplankton-dominated regions (North, West, Center, South). All samples were collected from open-lake areas, excluding river-inflow sites. The macrophyte-dominated East region was not surveyed. A total of 116 composite samples were collected during spring (1–3 April 2017) and summer (1–3 August 2017). Only representative sites are shown; full coordinates are listed in Table S1.

**Figure S2.**
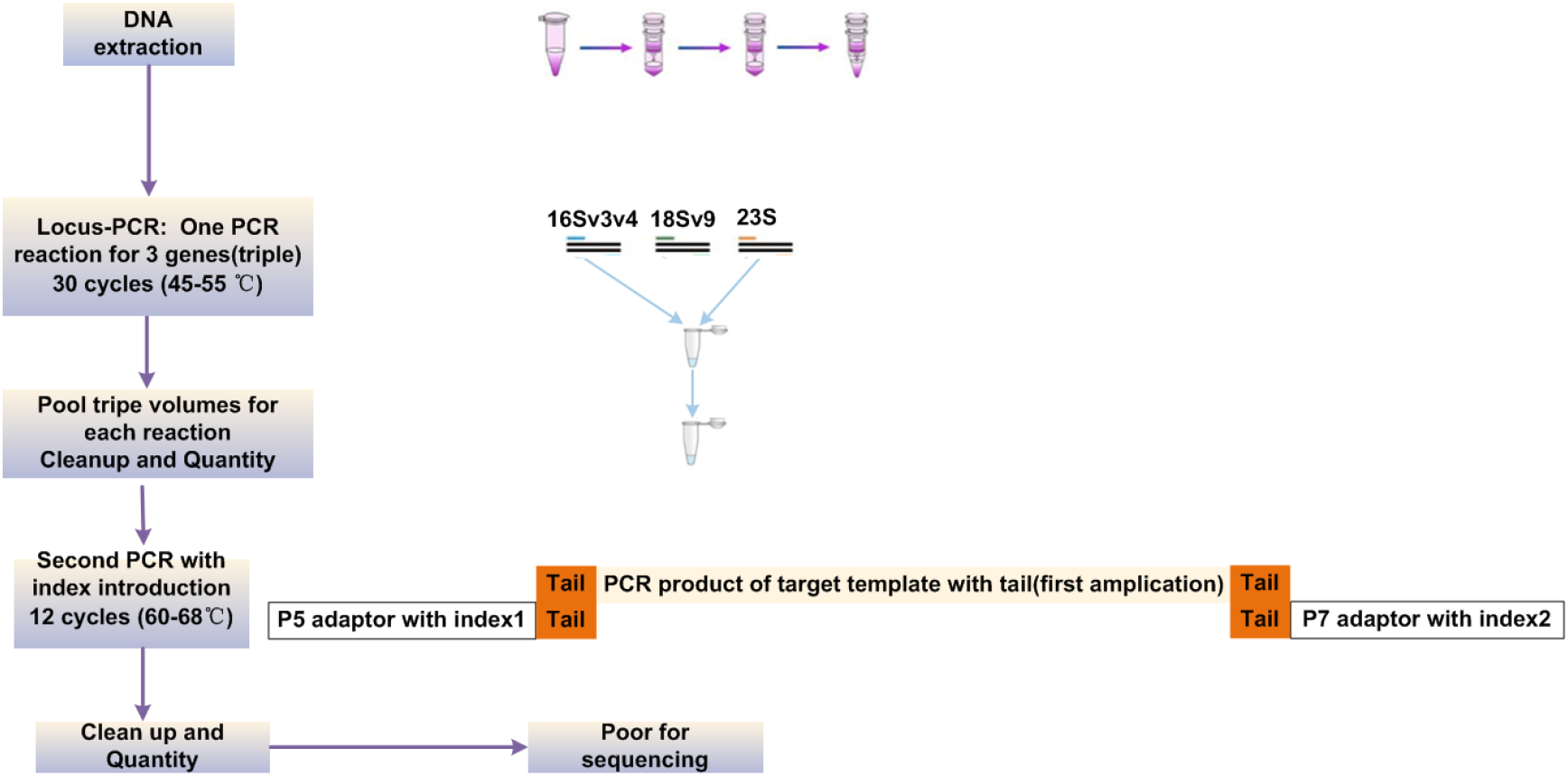
Workflow of multiplex library preparation and sequencing. Three loci (16S, 18S, 23S) were co-amplified with TruSeq-tailed primers, pooled, purified, and sequenced on Illumina MiSeq (2 × 300 bp).

**Figure S3.**
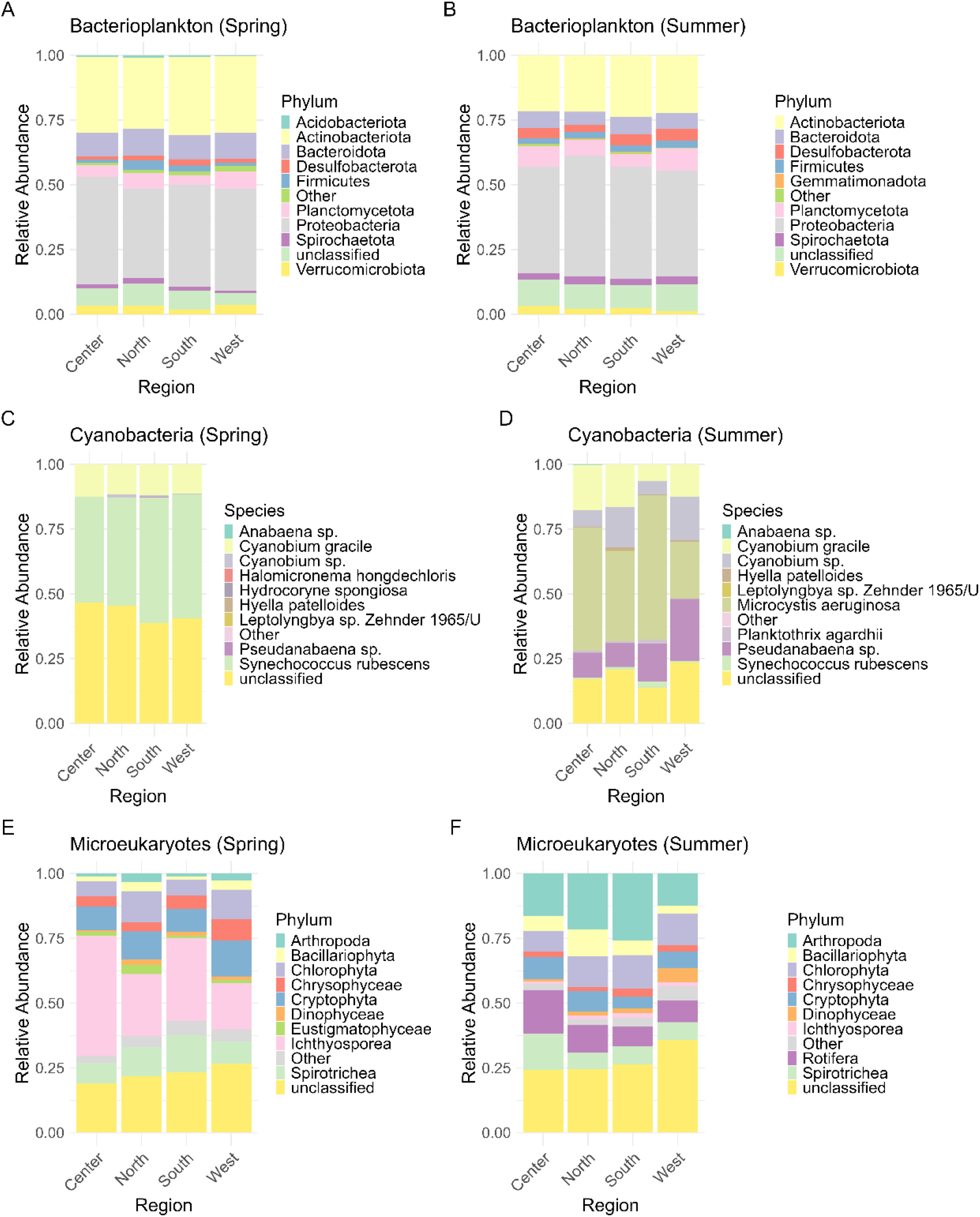
Seasonal and regional composition of bacterioplankton, cyanobacteria, and microeukaryotes. Stacked barplots show top 10 taxa per season; remaining taxa pooled as “Other.”. Relative abundances were normalized within each region..

**Figure S4.**
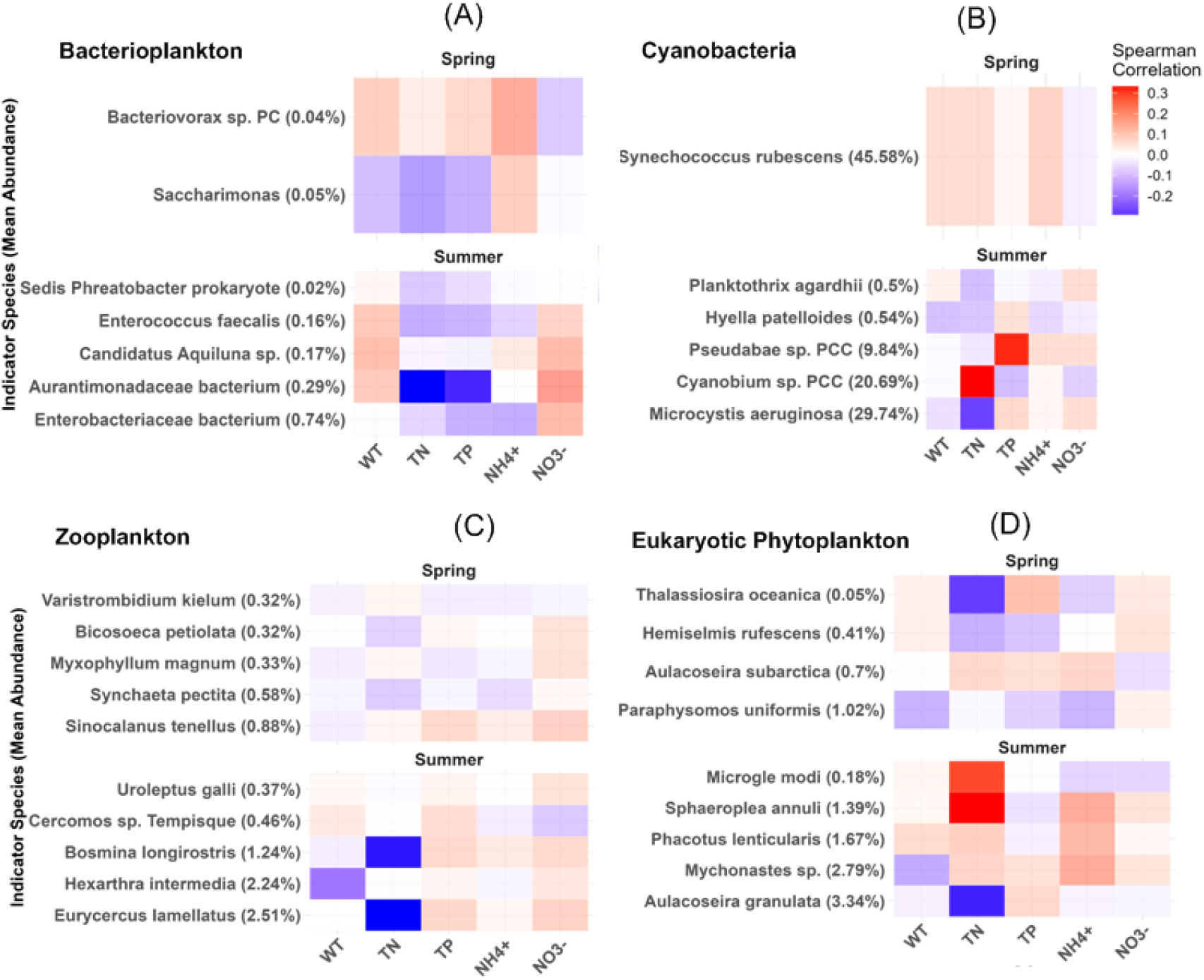
Spearman correlations between indicator taxa and environmental variables for (A) bacterioplankton, (B) cyanobacteria, (C) zooplankton and (D) eukaryotic phytoplankton. Indicator taxa identified by IndVal (p < 0.01); when none detected, top 5 taxa were used. Red/blue denote positive/negative correlations; significant cells (p < 0.05) shown at full opacity.

**Figure S5.**
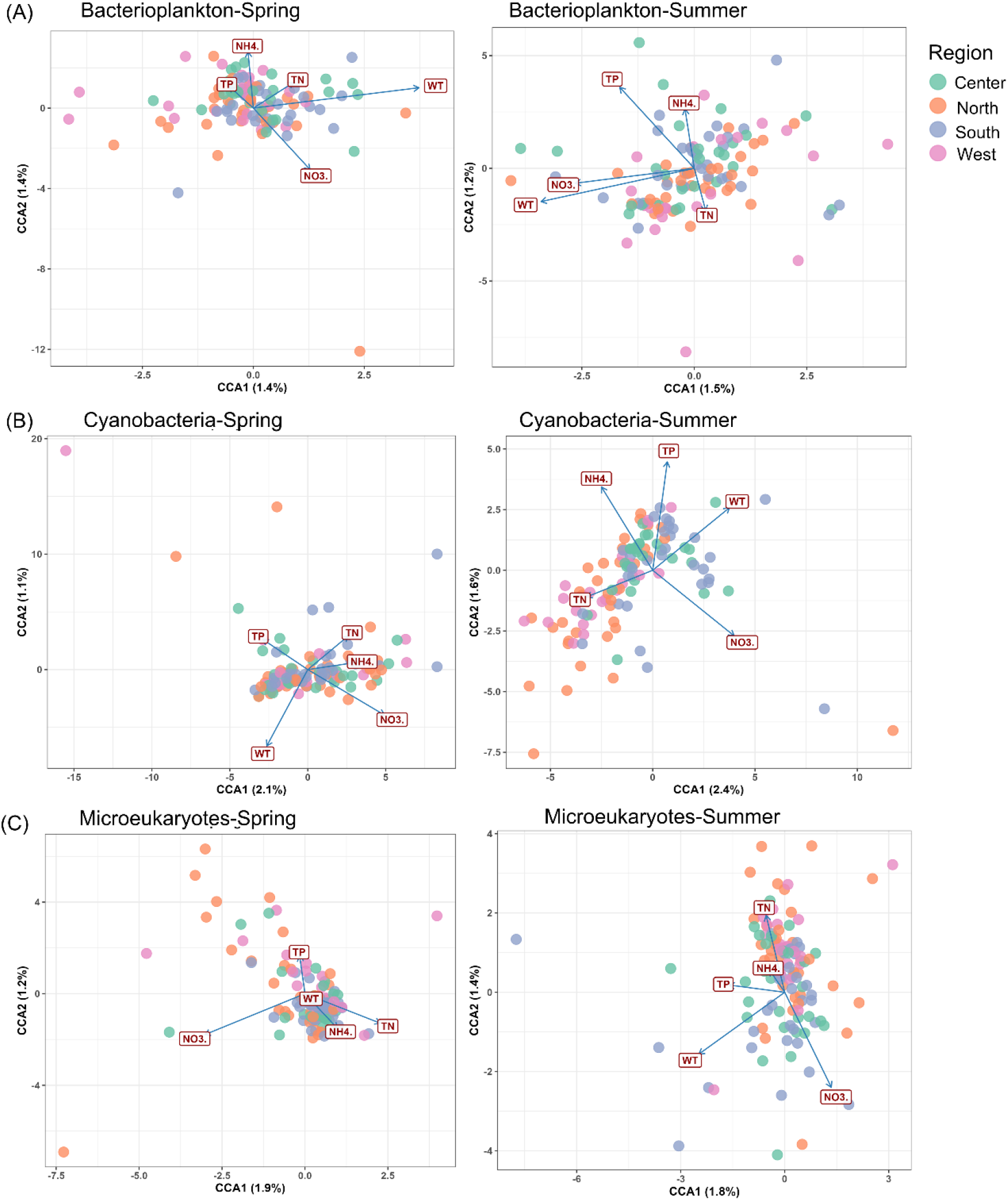
CCA of microbial communities for bacterioplankton (A), cyanobacteria (B), and microeukaryotes (C). Points colored by region; arrows show significant environmental gradients. Variance explained by the first two axes is indicated. WT, water temperature; TN, total nitrogen; TP, total phosphorus; NH₄⁺-N and NO₃⁻-N shown as “NH4.” and “NO3.”.

**Figure S6.**
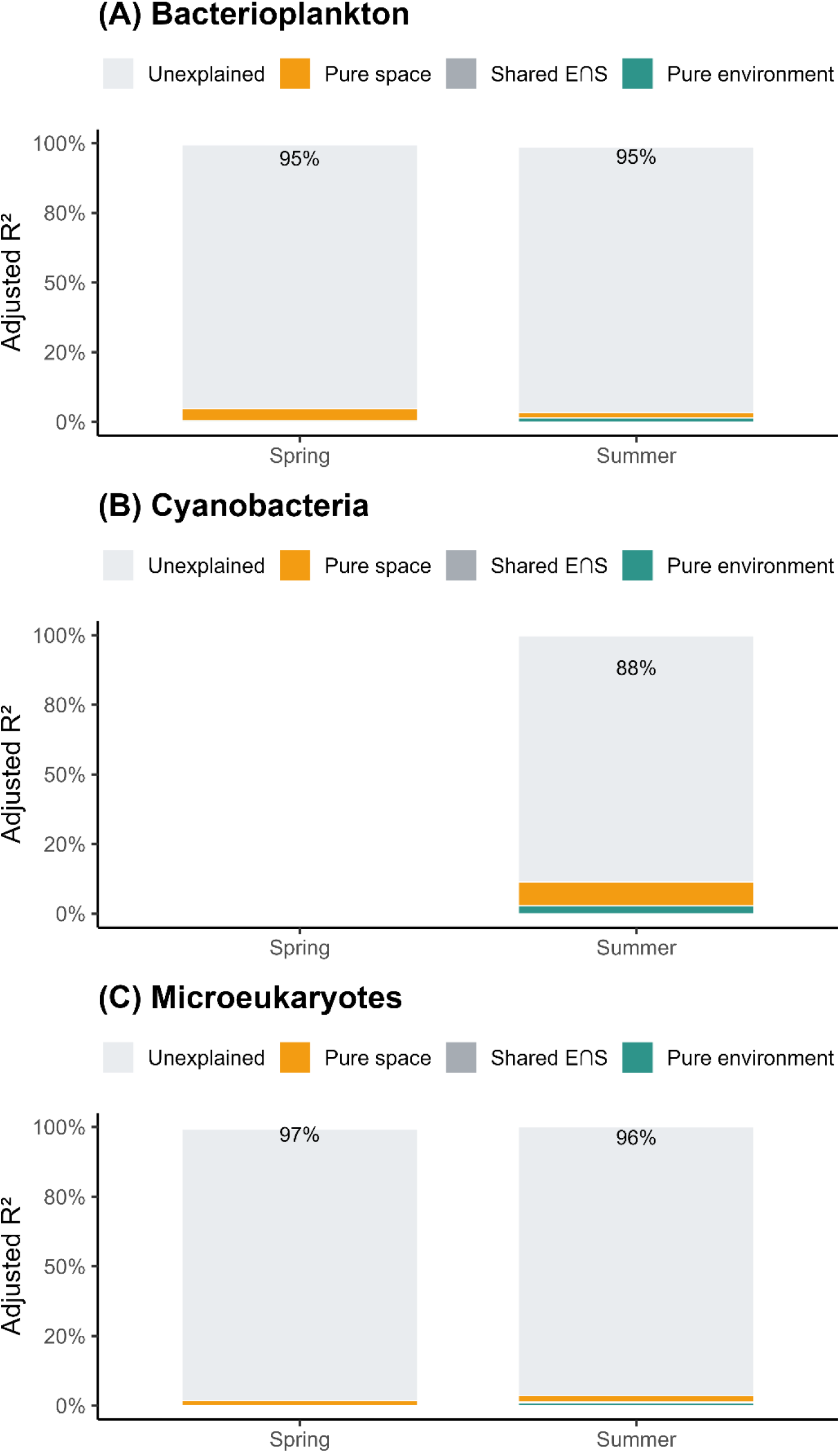
Seasonal variation partitioning into environmental, spatial/dbMEM, shared, and unexplained fractions for bacterioplankton (A), cyanobacteria (B), and microeukaryotes (C). Percentages in “Unexplained” indicate variance not accounted for by environment or space.

**Figure S7.**
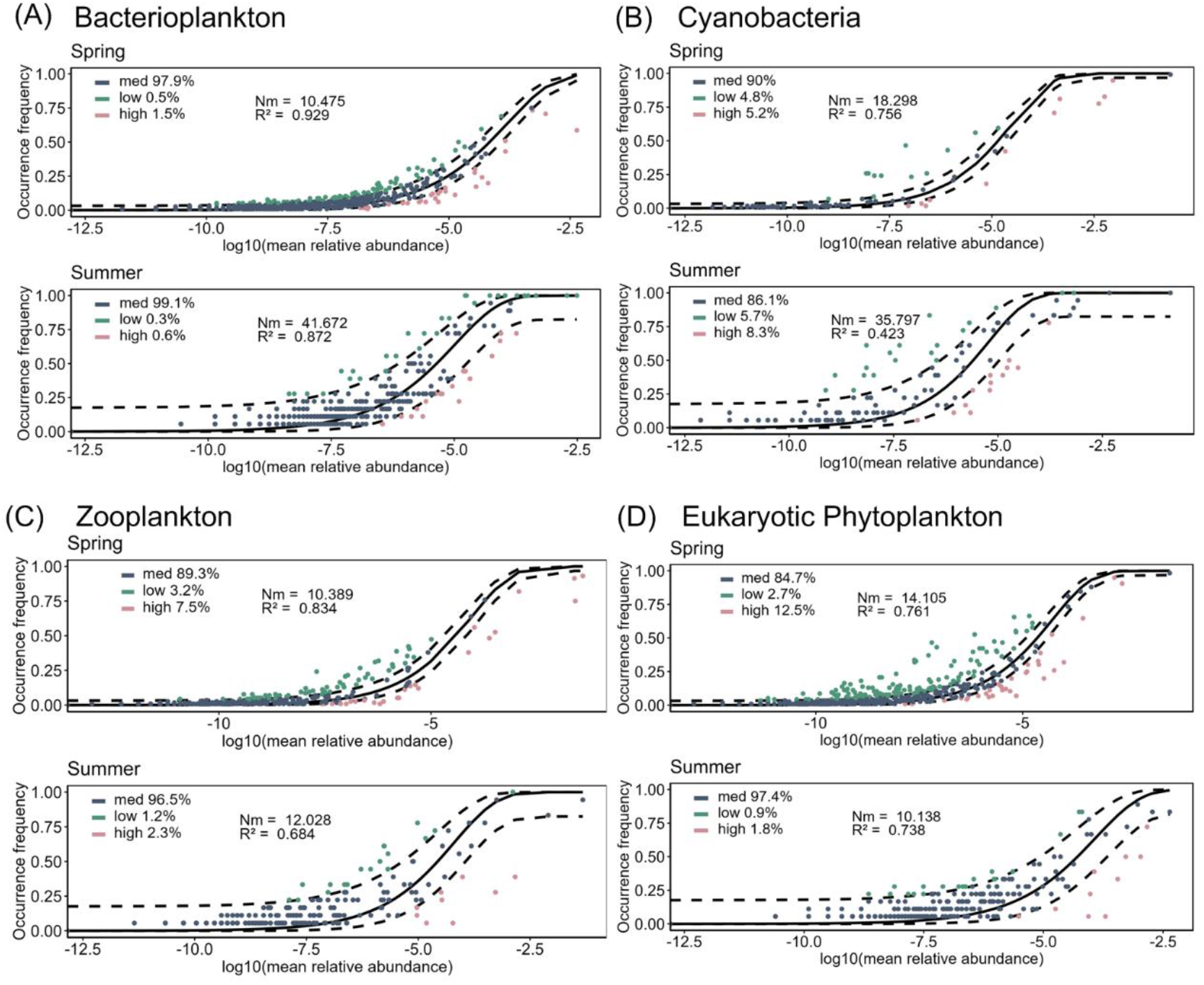
Neutral community model fits for spring and summer of (A) bacterioplankton, (B) cyanobacteria, (C) zooplankton and (D) eukaryotic phytoplankton. Solid lines show best-fit models; dashed lines show 95% CIs. ASVs classified as neutral, less frequent, or more frequent than predicted. R² and Nm shown.

**Figure S8-10.**
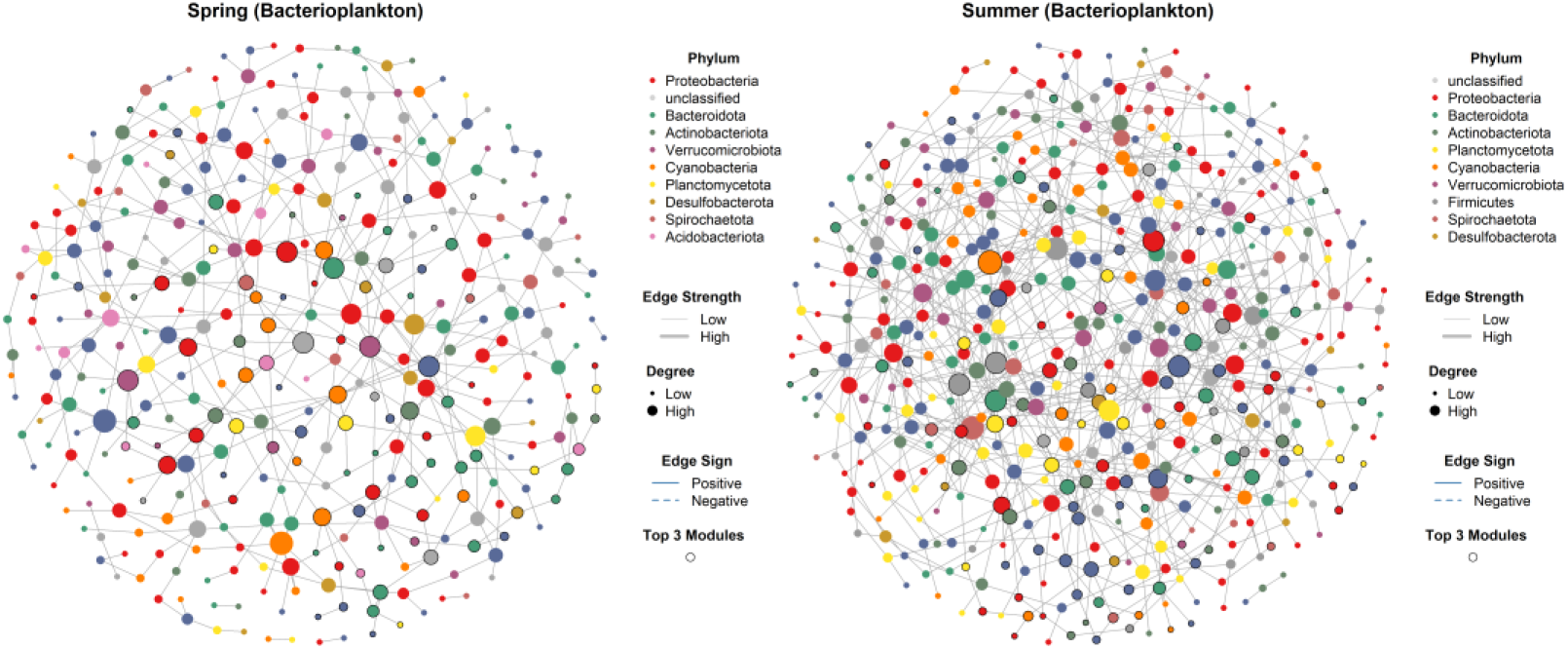
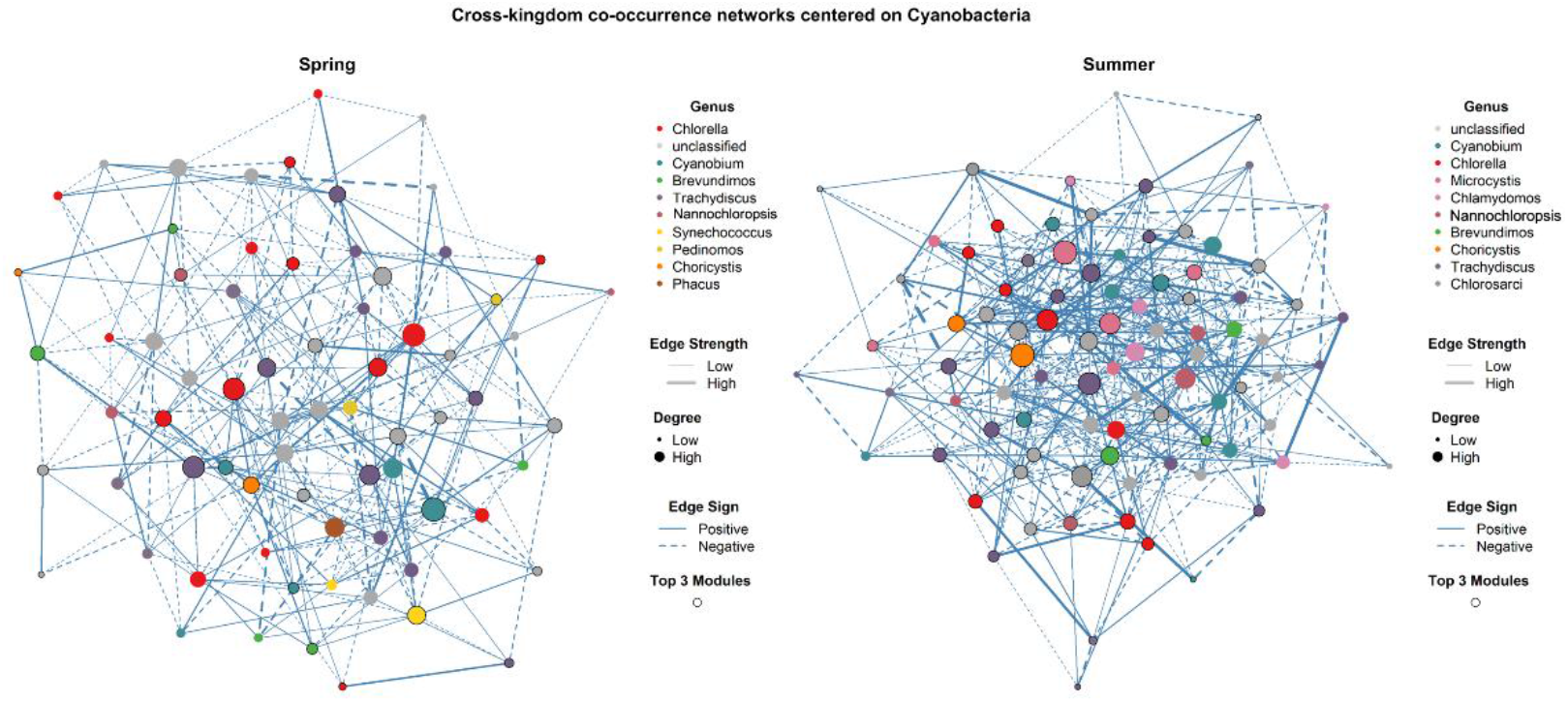
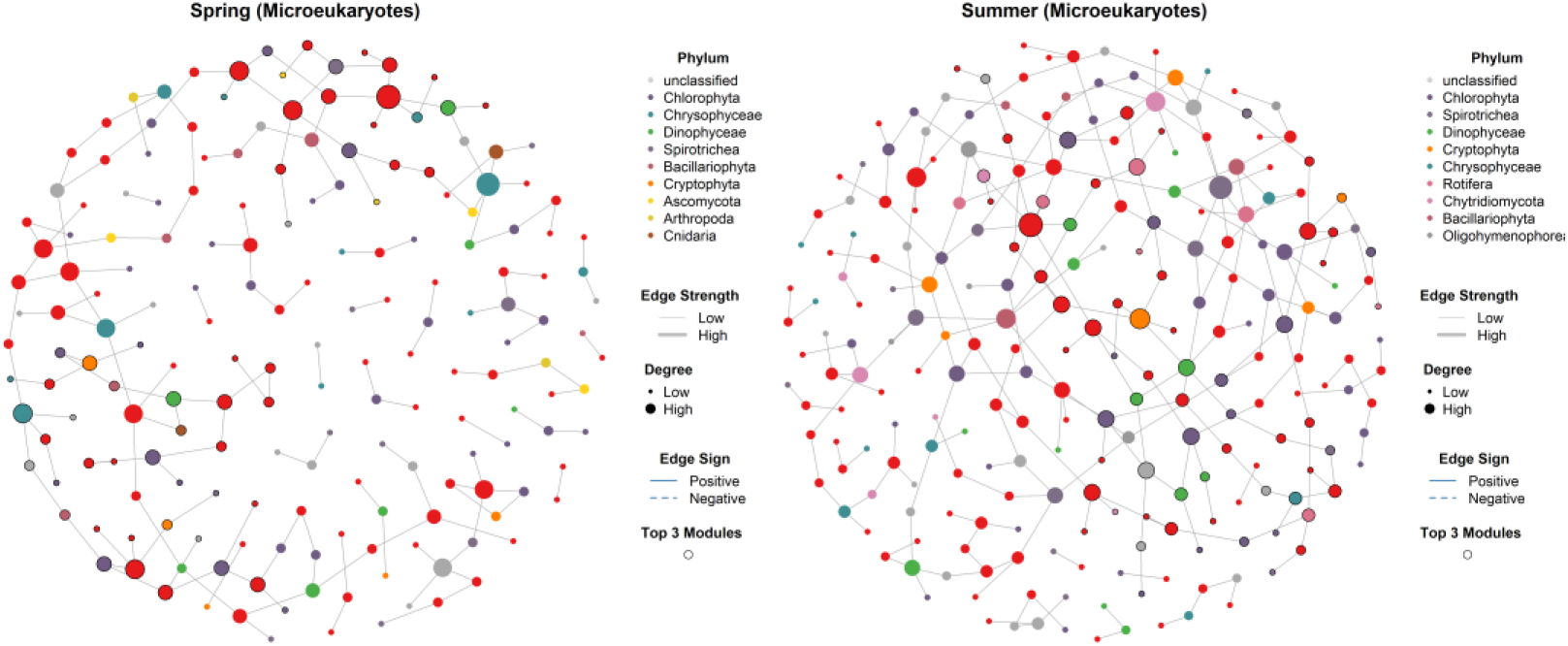
Spring and summer SparCC networks for bacterioplankton (S8), cyanobacteria (S9), and microeukaryotes (S10). Node colors indicate taxonomy; node sizes show degree. Solid/dashed edges represent positive/negative correlations. Top modules are outlined. For S9, the network includes all taxa co-occurring with cyanobacterial ASVs to capture cross-kingdom interactions, leveraging the 23S marker’s ability to simultaneously amplify cyanobacteria and eukaryotic plastids.

**Figure S11.**
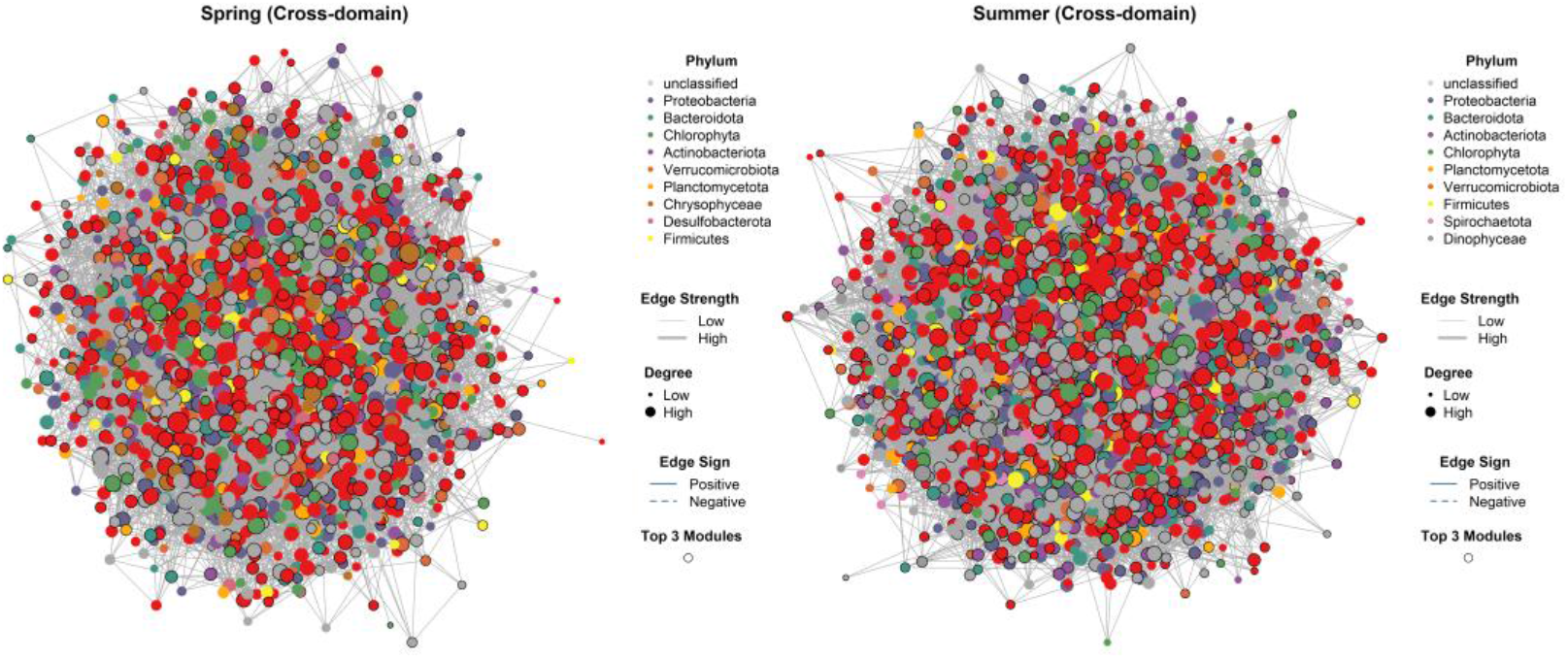
Cross-domain co-occurrence networks in spring (A) and summer (B). Bacteria and eukaryotes shown at the phylum level; cyanobacteria at the order level. Node colors reflect taxonomic identity; solid/dashed edges represent positive/negative associations; thick edges highlight cross-group links.

**Figure S12.**
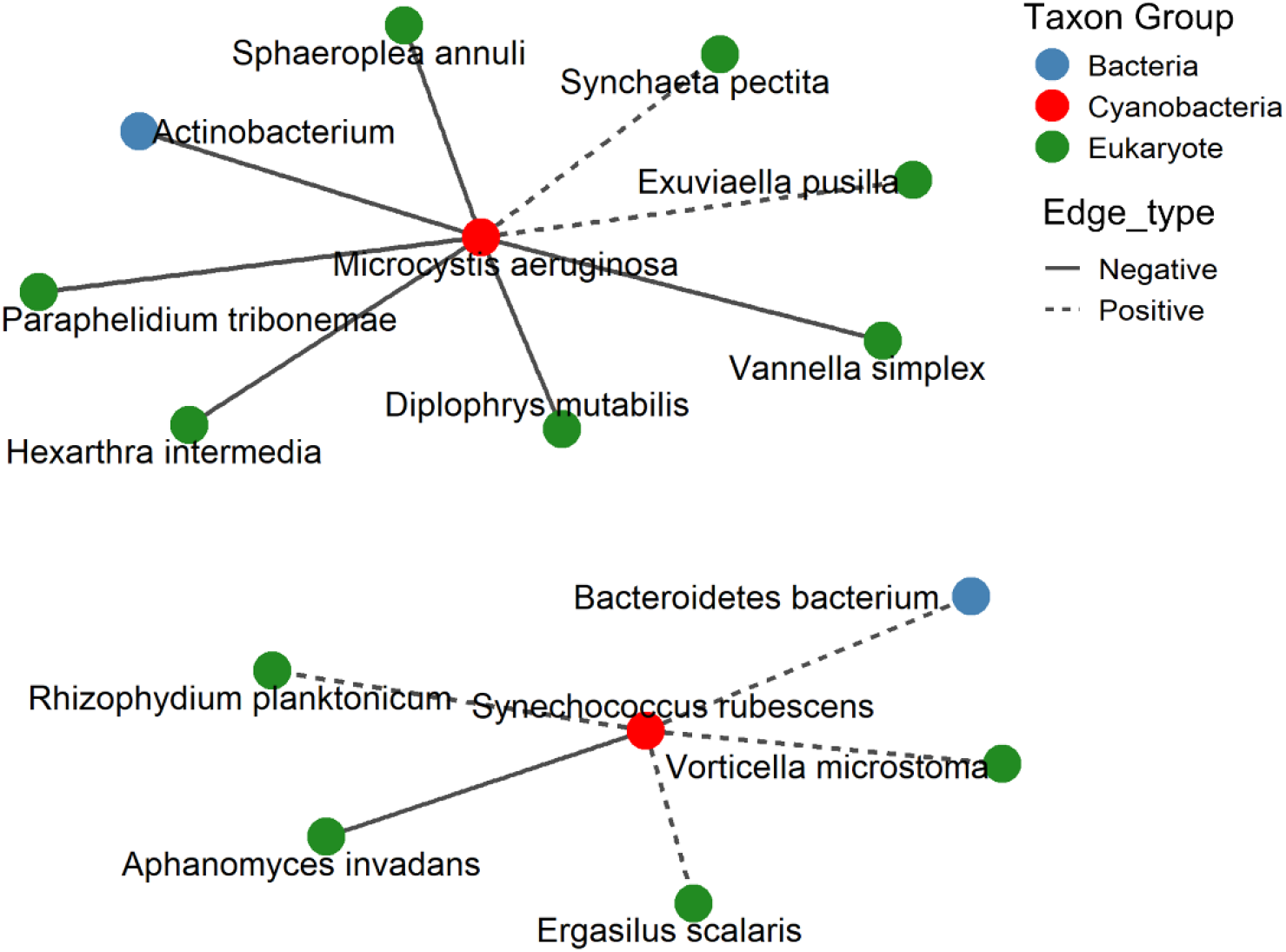
Species-level networks showing interactions of *Synechococcus rubescens* (spring) and *Microcystis aeruginosa* (summer). Nodes colored by domain; solid/dashed edges represent positive/negative correlations.

**Figure S13.**
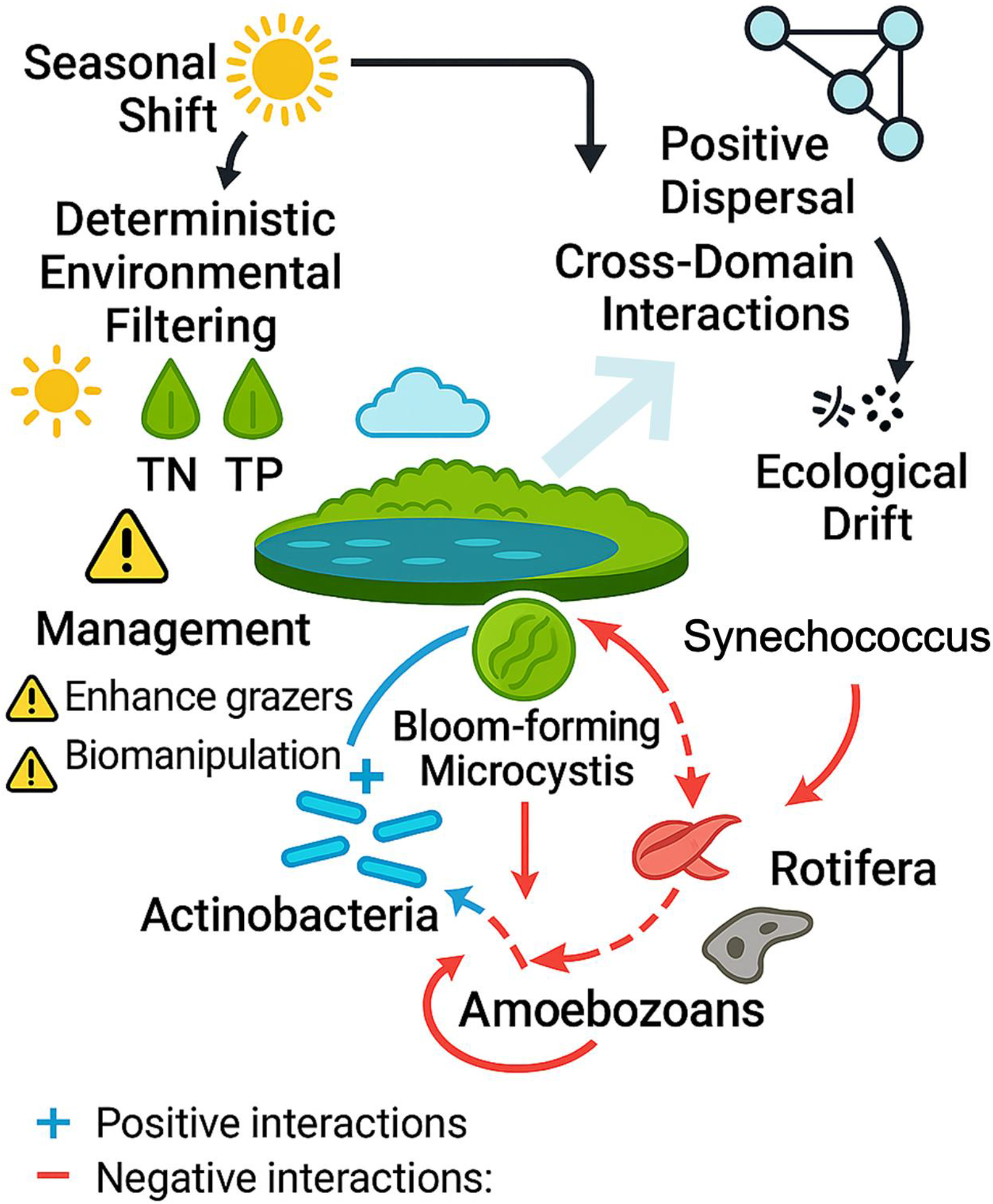
Conceptual diagram illustrating how seasonal transitions and cross-domain interactions influence *Microcystis* bloom development and highlighting potential intervention targets.

**Table S1.** Metadata and sequencing summary for all samples included in the multi-marker eDNA analysis.

**Table S2.** Multiplex PCR system and primers used in this study.

**Table S3.** Taxonomic assignments of three genetic markers for multiple microbial groups.

**Table S4.** Total taxonomic assignments of multiple microbial groups at both seasons. **Table S5.** Summary of PERMANOVA tests evaluating the effects of Season, Region, and Season × Region on multiple microbial community β-diversity.

